# Highly synchronized cortical circuit dynamics mediate spontaneous pain

**DOI:** 10.1101/2022.10.04.510722

**Authors:** Weihua Ding, Lukas Fischer, Qian Chen, Ziyi Li, Liuyue Yang, Zerong You, Kun Hu, Xinbo Wu, Xue Zhou, Wei Chao, Peter Hu, Tewodros Mulugeta Dagnew, Daniel Dubreuil, Shiyu Wang, Suyun Xia, Caroline Bao, Shengmei Zhu, Lucy Chen, Changning Wang, Brian Wainger, Peng Jin, Jianren Mao, Guoping Feng, Mark T. Harnett, Shiqian Shen

## Abstract

Cortical neural dynamics mediate information processing for the cerebral cortex, implicated in fundamental biological processes, such as vision and olfaction, in addition to neurological and psychiatric diseases. Spontaneous pain is a key feature of human neuropathic pain. Whether spontaneous pain pushes cortical network into an aberrant state, and if so, whether it can be brought back to a ‘normal’ operating range to ameliorate pain are unknown. Using a clinically relevant mouse model of neuropathic pain with robust spontaneous pain-like behavior, we report that orofacial spontaneous pain activated a specific area within the primary somatosensory cortex (S1), displaying synchronized neural dynamics revealed by intravital two-photon calcium imaging. This synchronization was underpinned by local GABAergic interneuron hypoactivity. Pain-induced cortical synchronization could be attenuated by manipulating local S1 networks or clinically effective pain therapies. Specifically, both chemogenetic inhibition of pain-related c-Fos-expressing neurons, and selective activation of GABAergic interneurons, significantly attenuated S1 synchronization. Clinically effective pain therapies including carbamazepine and nerve root decompression could also dampen S1 synchronization. More importantly, restoring a ‘normal’ range of neural dynamics, through attenuating pain-induced S1 synchronization, alleviated pain-like behavior. These results suggest spontaneous pain pushes S1 regional network into a synchronized state, whereas reversal of this synchronization alleviates pain.

## Introduction

While the functions of cortical circuits vary dramatically between regions and species, general principles have been found which constrain neural dynamics to stable activity regimes. For example, sleep, quiet wakefulness, and locomotion all exhibit characteristic global electrophysiological signatures that originate from widely varying, but still healthy, states of synchrony^1–6^. Within these states, cortical dynamics are broadly stable, so that finely tuned synaptic connectivity adjustments can produce adaptive changes in behavioral output.

Substantial experimental^7, 8^ and computational^9^ work has delineated a number of mechanisms that cortical circuits use to stabilize their activity. A precise coordination of excitation and inhibition is the key component of this dynamical stability, implemented by processes including local circuit wiring rules^10^, homeostatic regulation of intrinsic excitability^11^, and synaptic scaling^12^. It is well-known that disease states exhibit abnormal cortical dynamics^4^, but it is not clear how, or even if, this dysregulated activity actually drives maladaptive behavior. It is additionally unknown what specific cellular or circuit mechanisms produce altered cortical dynamics in disease states. While ectopic positive feedback loops have been implicated in epileptic seizures^13^ and depression^14^, it is unknown if this is a general mechanism. Although evidence points to alterations in the dynamical stability mechanisms mentioned above, the connection(s) between disease states and aberrant cortical mechanisms has been challenging to study, given the lack of effective translational models for many human disorders.

Chronic neuropathic pain affects 11-40% of all adults^15^ and can be a devastating and life altering disease. In a large human study of chronic neuropathic pain, less than one third of patients report evoked pain, including mechanical hyperalgesia and allodynia^16^. Spontaneous pain, which occurs in the absence of external stimuli, represents a key aspect of human neuropathic pain with poorly understood etiology. Interestingly, anti-seizure medicine such as carbamazepine is a first line treatment for some chronic neuropathic pain conditions, such as trigeminal neuralgia (TN), suggesting that abnormal, hypersynchronous cortical activity may contribute to this disease state. Indeed, recent studies have reported an increase in correlation between cortical neurons in sensory cortex in evoked pain^17^. This suggests that the pain state may entrain a fraction of cortical neurons, potentially leading to abnormally hyper-synchronous patterns of activity that pushes cortical dynamics outside of their normal operating range.

Like many other human brain disorders, investigating the neural dynamics underlying chronic neuropathic pain has been difficult due to technical challenges of reliably inducing and parametrizing animal models that reflect the human pain experience, particularly spontaneous pain. Existing studies have investigated synaptic plasticity and cortical activity patterns in evoked pain with a particular focus on the anterior cingulate cortex^18^. Additionally, recent evidence suggests that pain may also alter neural dynamics in the primary sensory cortex (S1)^17, 19^. How spontaneous pain alters neural activity patterns in S1, and how these patterns may be brought back into normal operating range to ameliorate pain, are unknown.

We developed a novel pain model for inducing neuropathic pain via a simple impingement of the trigeminal nerve root. Mice that have undergone this procedure show hallmarks of TN, including spontaneous bouts of pain attacks, allodynia, avoidance of chewing solid food, and a range of other phenotypes that are also observed in humans. Using two-photon calcium imaging we directly observed how cortical neural dynamics in S1 are altered by pain. Additionally, we found that this cortical synchronization was underpinned by local GABAergic interneuron hypoactivity. Pain-induced cortical synchronization could be attenuated by manipulating S1 networks or clinically effective pain therapies. Attenuation of S1 synchronization reliably alleviated pain-like behavior. Together, our results provide new mechanistic insights into how neuropathic pain induces abnormal activity states in the cortex that can lead to debilitating pain behaviors including spontaneous pain for the subject.

## Results

### Robust neuronal activation observed in the primary somatosensory cortex (S1)

In the somatosensory homunculus, orofacial area represents the largest projection within the S1, among all body areas^20, 21^. We therefore reasoned that an orofacial neuropathic pain model with robust spontaneous pain - like behavior would facilitate mechanistic query of cortical substrates of pain. Human trigeminal neuralgia (TN) is a prototypic neuropathic pain condition characterized by lancinating pain, including both evoked pain and spontaneous paroxysmal pain attacks. Compelling clinical evidence indicates that trigeminal nerve root compression is one of its major causes^22^. We therefore modeled TN by mimicking the human pathology of trigeminal nerve root impingement at its entry zone^23^. We discovered that a natural orifice, the foramen lacerum, lies underneath the trigeminal nerve root, across species (**Supplementary Figure 1a**). Taking advantage of this, we applied Surgifoam to create trigeminal nerve root compression (Foramen Lacerum Impingement of Trigeminal nerve root, FLIT) (**Figure 1a and Supplementary Figure 1b**). Besides mechanical allodynia (**Figure 1b**), the FLIT model demonstrated robust spontaneous behavior that was likely related to pain, including excessive spontaneous facial grooming (**Supplementary Figure 1c, movie S1**), and spontaneous paroxysmal asymmetrical facial grimacing with intermittent eye squinting, ipsilateral to the side of nerve compression (**Figure 1c; Supplementary Figure 1d; movie S2**). Uncontrollable facial twitching or tic-like grimacing is commonly seen in TN patients; facial grimacing has been increasingly accepted as an indication of pain in animals^24^. Spontaneous paroxysmal asymmetrical facial grimacing, therefore, may represent paroxysmal pain attacks, a key feature of human TN^23^. Moreover, we assessed the functional implications of pain, including body weight gain (**Supplementary Figure 1e**), wood chewing (**Supplementary Figure 1f**), incisors length (**Supplementary Figure 1g**) and solid vs. soft chew preference (**Supplementary Figure 1h to j**). Compared with Sham control, the FLIT model was associated with an initial phase of weight loss followed by significantly less body weight gain; incisors overgrowth; minimal wood chewing; and avoidance of solid chew. As the trigeminal nerve is a mixed nerve and its motor innervation controls masseter muscles, we ruled out masseter muscle atrophy as the cause of pain-like behavior (**Supplementary Figure 2a to i**). For a global assessment of pain, we computed composite pain scores consisting of six pain - related behaviors, including mechanical withdrawal thresholds, spontaneous grooming, body weight change, incisors length increase, food preference, and wood chewing activity (see Methods), which provides a comprehensive assessment of evoked pain, spontaneous behavior, as well as functional implication of pain (**Figure 1d**). Moreover, we found the FLIT model was associated with anxiety - like behavior (**Supplementary Figure 3a to c**) and sexual dysfunction (**Supplementary Figure 3d to f**), further substantiating its clinical relevance.

**Figure 1.**
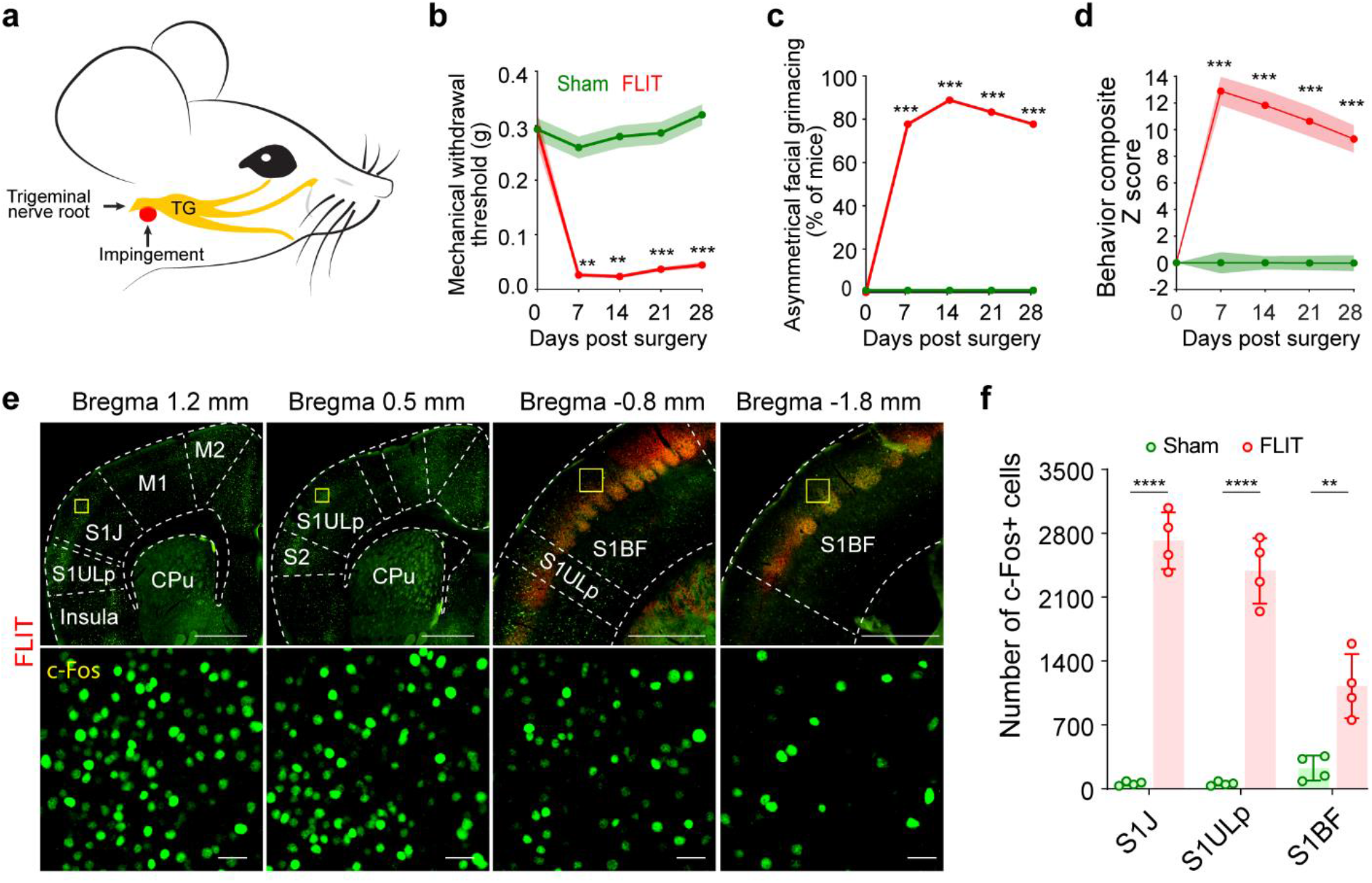
FLIT model of trigeminal neuralgia leads to robust c-Fos activation in S1ULp-S1J. **(a)** Diagram of trigeminal nerve root impingement to recapitulate human TN. Yellow structure depicts trigeminal anatomy including trigeminal nerve root, trigeminal ganglion and peripheral branches; Red represents Surgifoam impingement site at the trigeminal nerve root. **(b to d)** Behavioral testing for the FLIT model. Mice underwent Sham (*n* = 18) and FLIT (*n* = 18) surgery, followed by behavioral testing at indicated time points. **(b)** Mechanical withdrawal threshold to von Frey filament (mean ± SEM). Two-way ANOVA with Bonferroni post-hoc testing indicates significant differences between groups (\*\**P* < 0.01; \*\*\**P* < 0.001). **(c)** Percentage of mice with asymmetrical facial grimaces. Mice in FLIT group displayed paroxysmal asymmetrical facial grimaces. Fisher’s exact test was used to determine statistical difference (\*\*\**P* < 0.001). **(d)** Summary quantification of behavioral tests quantified as composite Z score (mechanical withdrawal; grooming; body weight; incisors length; wood weight changes; percentage of time eating solid chew) was computed over 28 days. Two-way ANOVA with Bonferroni post-hoc testing indicates statistic difference between groups (\*\*\**P* < 0.001). **(e)** c-Fos activation in S1ULp-S1J after surgery. Representative tangential slices of c-Fos staining of FLIT mice at 4X and 10X. Sequential slices from left to right represent coronal sections covering S1J (bregma 1.2 mm), S1ULp (bregma 0.5 mm), anterior S1BF (bregma −0.8 mm) and posterior S1BF cortex (bregma −1.8 mm). Slices between bregma −0.8 mm and −1.8 mm were co-stained with VGLUT2 (red) to visualize barrels. Lower panels represent boxed regions of corresponding upper panels. **(f)** Quantification of c-Fos positive cells in the S1J, S1ULp and S1BF cortex (*n* = 4 per group). Two-way ANOVA with Bonferroni post-hoc testing indicates significant difference between the groups (\*\**P* < 0.01, \*\*\*\**p* < 0.0001).

To examine the cortical activation in response to pain, c-Fos was immunostained in tangential slices of the brain two-hour after FLIT or Sham surgery (**Supplementary Figure 4a and b**). These mice did not undergo any mechanical or thermal testing to minimize behavioral test - evoked pain or c-Fos expression. VGLUT2 was used as a co-stain to identify the barrel cortex (BF). Brain regions including M1 (primary motor cortex), M2 (secondary motor cortex), insula, PFC (prefrontal cortex), S1, and S2 (secondary somatosensory cortex) were compared between the FLIT and Sham controls. Among these regions, S1 displayed the most striking increase in c-Fos^+^ cells (**Figure 1e, Supplementary Figure 4c**). More importantly, S1ULp (upper lip)- S1J (jaw) exhibited the most dramatic increase among all cortical regions examined. Interestingly, S1BF, bordering S1 ULp-S1J, exhibited only moderate increases in c-Fos^+^ cells (**Figure 1f**), and these cells were primarily located at the anterolateral of BF bordering S1ULp. When VGLUT1 and GAD67 were used to co-stain for excitatory and inhibitory neurons, respectively, most c-Fos^+^ cells were VGLUT1^+^, indicating majority of these cells were excitatory neurons (**Supplementary Figure 4d and e**). The FLIT model, therefore, exhibited a behavioral battery with robust features of spontaneous pain which was accompanied by distinct c-Fos expression in the S1ULp -S1J region, consistent with cortical activation.

### Cortical synchronization in the S1ULp – S1J region induced by pain

To directly assess the cortical neural dynamics associated with pain, AAV-CaMKII-GCaMP6f was microinjected to a relatively wide cortical area (2-3 mm X 3-3.4 mm) of C57BL/6 mice (**Figure 2a**). Animals then underwent Sham or FLIT surgery, followed by intravital two-photon imaging, with both lower magnification (4X, wide-field) and higher magnification (20X). Mice were head-fixed without using any anesthesia, as anesthetics are known to suppress cortical neural activities^25^. More importantly, animals were free of experimentally imposed stimuli to minimize evoked pain. At 7 days post FLIT surgery, under lower magnification, mice in the FLIT group displayed relatively higher neural activities compared with Sham group, consistent with previous reports of cortical activation in response to pain^26^,^27^. More interestingly, neural activities among the examined cortical areas (S1, S2, M1, M2, insula, etc) were not homogenous. Instead, they were significantly higher in a relatively small area within the S1 (**Figure 2b**), which corresponded to S1ULp-S1J, the same region with robust c-Fos expression shown in Figure 1. This region was then examined under higher magnification at single neuron resolution before and after induction of pain. Very surprisingly, after induction of pain, neurons in the S1ULp-S1J region started to fire synchronously in awake and resting state, in the absence of experimentally imposed stimuli. This synchronization pattern was absent in all animals before induction of pain or Sham group at all time points. Using 50% of imaged neurons with simultaneous activation as a cut-off, bouts of spontaneous synchronous firing occurred once every 2-5 minutes on average (**Figure 2c and d; Supplementary Figure 5a to c and h to j; movie S3).** For any imaging field, each neuron was computed against all other neurons to derive a correlation index for quantification of the synchronicity of neuronal firings. Results showed that the FLIT model had significantly higher correlation indices (**Figure 2e to h; Supplementary Figure 5d to g and k to m**). Additionally, composite scores of pain-like behavior and S1ULp-S1J neuronal synchronization indices were plotted for correlation analysis, and there was a statistically significant correlation between these two parameters (**Figure 2i**), suggesting a plausible link between neural synchronization and pain.

**Figure 2.**
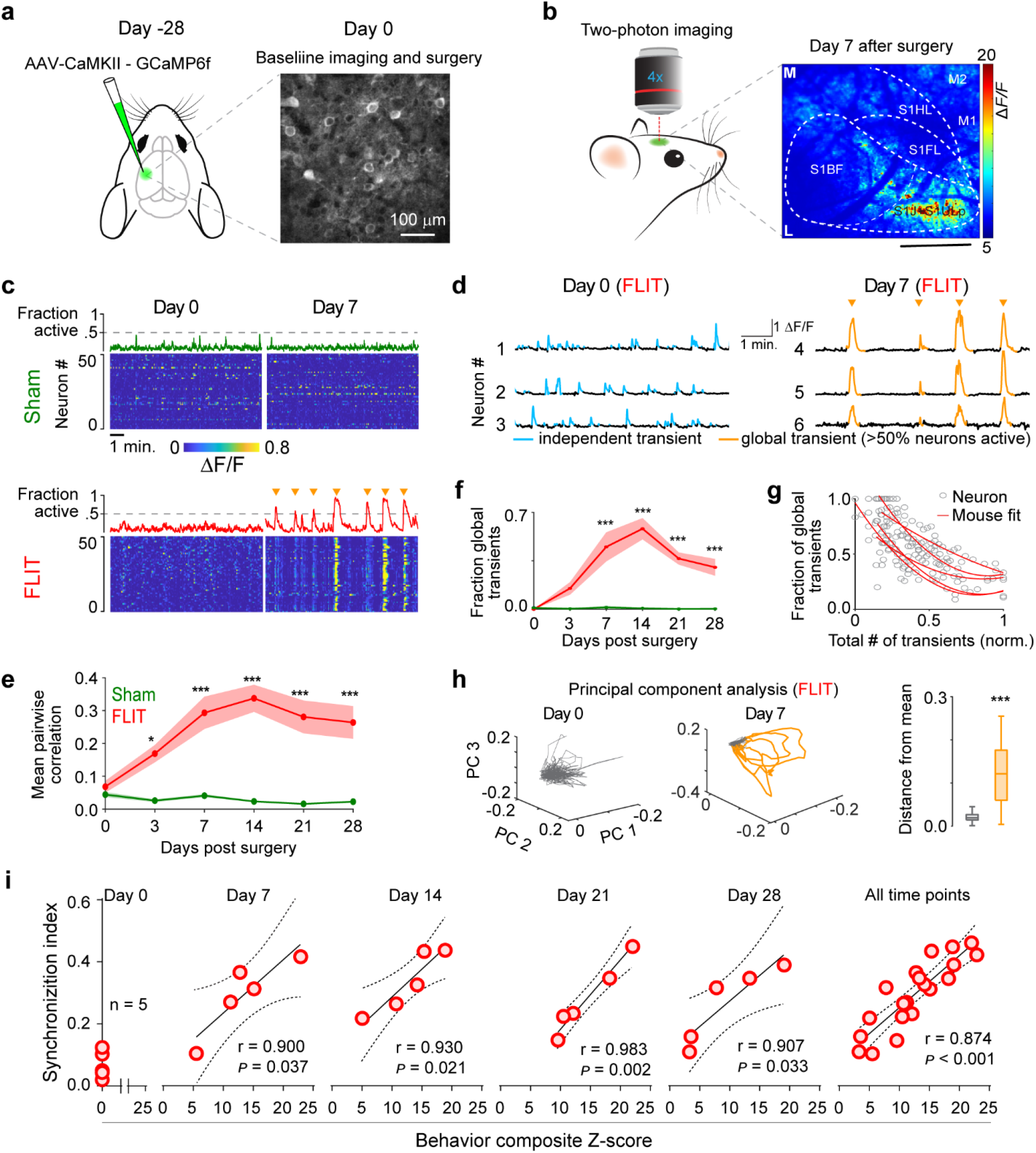
Highly synchronized S1 neural dynamics in FLIT mice. **(a to h)** Two-photon calcium imaging of anterolateral S1 cortex in awake mice at resting state (*n* = 7 Sham, *n* = 5 FLIT). Images were acquired from the same field of view across all days. **(a)** Left panel: schematic of AAV injection to S1 cortex contralateral to the FLIT surgery side. Right panel: GCaMP6 expression mediated by AAV8-CaMKII-GCaMP6f in S1 cortex. Shown representative picture was obtained under two-photon microscope at 20X. **(b)** Robust neuronal activities in S1ULp-S1J captured by widefield two-photon calcium imaging in large cortical areas of mice expressing AAV8-CaMKII-GCaMP6f. Heatmap of calcium activities of the imaging field seven days after FLIT surgery. M: Middle, L: lateral. **(c)** Representative heatmaps and corresponding fraction of simultaneously active neurons for each group at days 0 and 7. Orange triangles indicate global events (>50% of neurons active simultaneously). **(d)** Sample neuron calcium transients from S1 of a FLIT mouse at day 0 and day 7. Golden indicates global synchronized events. **(e)** Mean pairwise correlation co-efficient across days and groups. The Sham group did not exhibit significant correlation at any timepoints, whereas the FLIT group exhibited significantly higher correlation (Two-way ANOVA: *P* < 0.001 *vs*. Sham group, pairwise comparison using Bonferroni post hoc test, \**P* < 0.05; \*\*\**P* < 0.001). **(f)** Fraction global transients across days and groups. The FLIT group exhibited significantly global transients after day 3. (Two-way ANOVA: *P* < 0.001 *vs*. Sham group, global transients comparison using Bonferroni post hoc test, \*\*\**P* < 0.001). **(g)** Fraction of global transients per neuron (normalized to each animal). Each ring indicates one neuron, each line represents the fit for one animal from the FLIT group. Sham group is not shown due to lack of global events. **(h)** Left and middle panels: Representative plots of neuronal trajectories using the first three coefficients of principal component analysis (PCA) in FLIT model mice at days 0 and 7. Activity during global events is highlighted in golden color. Right: Euclidean distance between the mean of the first 3 coefficients and global events (golden) *vs*. non-global events (gray, Wilcoxon rank-sum test: \*\*\**P* < 0.001). **(i)** Correlation between Z score and synchronization index in FLIT model. Composite Z scores composed of 6 pain-related behaviors were plotted against synchronization indices for individual animals (*n* = 5, each red circle represents one animal). There was a positive correlation between Z score and synchronization index at indicated time points. Solid line represents linear regression, dash line represents 95% confidence interval.

### Pain-induced c-Fos expressing neurons are responsible for synchronized neural dynamics

To examine the neural dynamics of pain-induced c-Fos expressing neurons, Fos^2A-iCreER^ (TRAP2) mice, which express tamoxifen-inducible Cre recombinase under the control of Fos promoter/enhancer elements^28^, were used to drive Cre-dependent GCaMP6f expression for intravital two-photon calcium imaging. Specifically, TRAP2 mice were microinjected with AAV-DIO-hSyn-GCaMP6f in the S1ULp - S1J and then underwent FLIT or Sham surgery in 4 weeks. Within 5 minutes of surgery, tamoxifen was administered intraperitoneally. Single dose tamoxifen was able to reliably induce Cre for 4 - 6 hours^29^. From day 7 post tamoxifen injection, intravital two-photon imaging was performed for Layer 2/3 neurons in awake mice (**Figure 3a**) to capture the neural dynamics of c-Fos-expressing neurons. In mice received FLIT surgery, strikingly synchronized neuronal activities were again observed in the S1ULp - S1J region (**Figure 3b to d**), similar to that observed in Figure 2. Synchronization of pain-related c-Fos expressing neurons suggests that these neurons might be responsible for the overall S1ULp-S1J synchronization in pain. To test this, S1ULp-S1J region of TRAP2 mice were microinjected with Cre-dependent hSyn-Gi-mCherry (Gi DREADD) and AAV-CaMKII-GCaMP6f followed by FLIT surgery in 4 weeks (**Figure 3e**). At day 7 post FLIT surgery, robust S1 ULp-S1J synchronization was observed (**Figure 3f**), consistent with Figure 2. Tamoxifen was then administered to facilitate c-fos - induced Cre expression. At day 14 post FLIT surgery, C21 was given to activate Gi DREADD^30^, S1 ULp-S1J neuronal activities were imaged prior to and after C21 administration. Results showed that inhibiting these pain-induced c-Fos expressing neurons largely abrogated the S1ULp - S1J synchronization (**Figure 3g and h**). As such, these neurons were likely the source of synchronized S1ULp-S1J neural dynamics.

**Figure 3.**
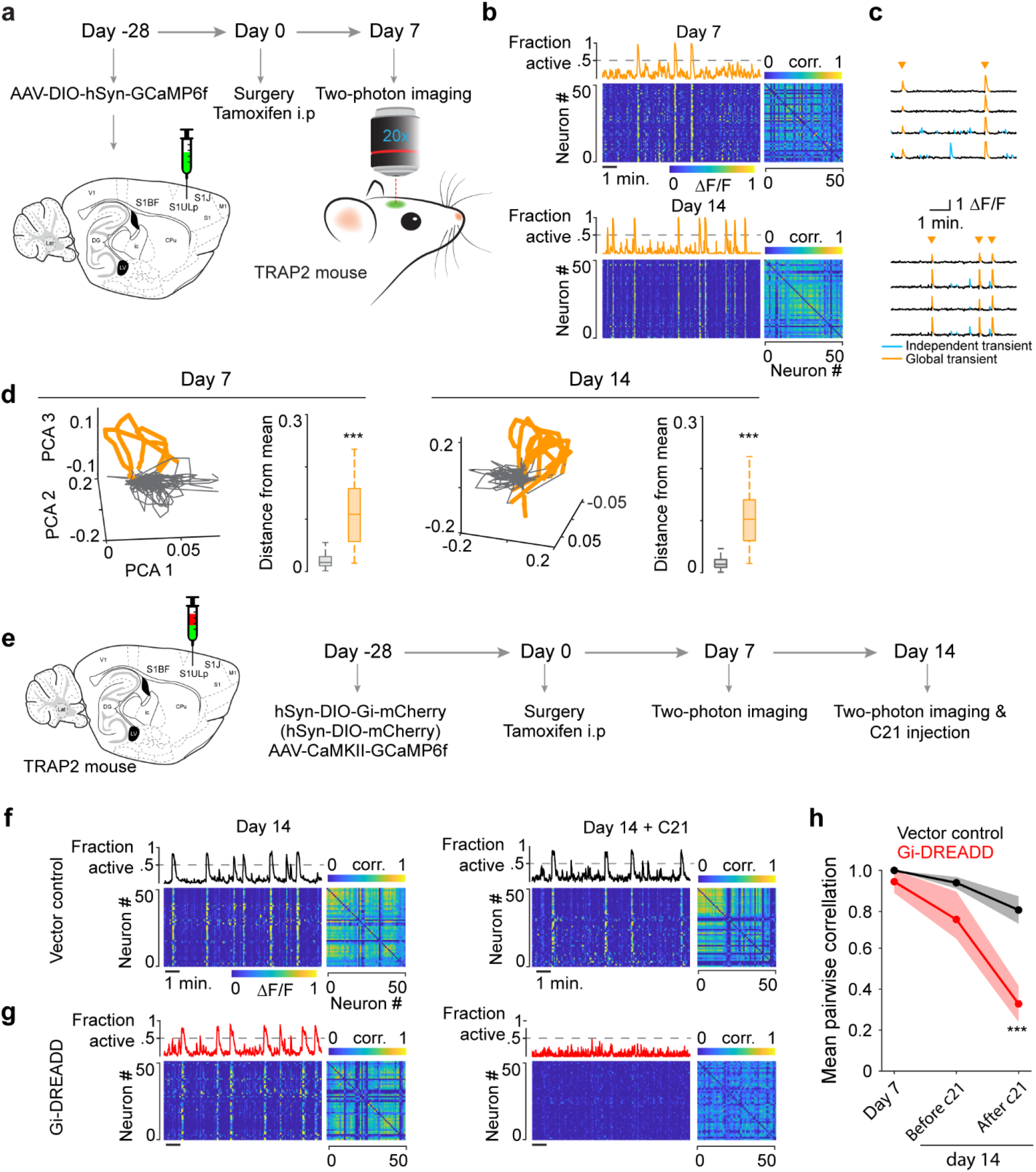
Pain-related c-Fos expressing neurons drives S1 synchronization. **(a to d)** Neuronal synchronization captured in S1ULp-S1J of TRAP2 mouse since day 7 after FLIT surgery and tamoxifen administration. **(a)** Diagram and flowchart of two-photon imaging in TRAP2 mouse. AAV-DIO-hSyn-GCaMP6f was injected in TRAP2 mouse 4 weeks ahead of FLIT surgery and tamoxifen administration. Intravital two-photon imaging was performed from day 7 after FLIT surgery and tamoxifen (*n* = 3). **(b)** Representative heatmaps with corresponding fraction of simultaneously active neurons and correlation matrices at day 7 and 14. Global synchronized neuron activity (>50% neurons active simultaneously) was presented in TRAP2 mice underwent FLIT surgery. **(c)** Sample neuron calcium transient traces from S1 of a FLIT mouse at day 7 and 14 after FLIT procedure. Golden arrowhead indicates global synchronized events (>50% of neurons simultaneously active). **(d)** Representative plots of neuronal trajectories using the first three coefficients of principal component analysis (PCA) in FLIT mice at days 7 and 14. Activity during global events is highlighted in golden color. Euclidean distance between the mean of the first 3 coefficients and global events (golden) *vs*. non-global events (gray, Wilcoxon rank-sum test: \*\*\**P* < 0.001). **(e to h)** S1ULp-S1J neuronal synchronization was subdued by inhibition of c-Fos induced Gi-DREADD expressing neurons in TRAP2 mice with TN. **(e)** Diagram and flowchart of inhibition of c-Fos induced Gi-DREADD expressing neurons in S1ULp-S1J of TRAP2 mouse. Mixed AAV-DIO-Gi-DREAD and AAV-CaMKII-GCaMP6f was injected in S1ULp-S1J of TRAP2 mouse 4 weeks ahead of FLIT surgery and tamoxifen administration. Two-photon imaging was performed before and after C21 at day 14 (*n* = 3). **(f)** Representative heatmap and correlation matrix showing neuronal synchronization presented since day 7 after FLIT and Tamoxifen. C21 administration in mice injected with vector virus did not alter synchronization. **(g)** Representative heatmap and correlation matrix showing neuronal synchronization presented since day 7 after FLIT and Tamoxifen. C21 administration in mice injected with Gi-DREADD virus suppressed neuronal synchronization. **(h)** Mean pairwise correlation co-efficient across days and groups. Both groups of mice injected vector and Gi-DREADD virus exhibited significant correlation since day 7 after FLIT surgery, whereas C21 administration decreased correlation in mice injected with Gi-DREADD compared with mice injected with vector virus. Two-way ANOVA followed by Bonferroni post hoc test to determine the difference of vector control vs. Gi-DREADD. \*\*\**P* < 0.001.

### S1ULp-S1J c-Fos expressing neurons mediate pain-like behaviors

To interrogate if the c-Fos expressing neurons were implicated in pain-like behaviors, TRAP2 mice underwent microinjection of Cre-dependent AAV-DIO-hSyn-Gi-mCherry (Gi DREADD) or AAV-DIO-hSyn-mCherry (vector control) in S1ULp-S1J followed by FLIT surgery in 4 weeks (**Figure 4a**). Tamoxifen was administered immediately after FLIT surgery to facilitate the induction of Cre expression. Seven days were allowed for Gi DREADD expression. At day 7, DREADD actuator drug, C21^30^, was injected intraperitoneally at 0.8 mg/kg twice daily, to assess the effects of inhibiting these c-Fos expressing neurons. Pain behaviors including paroxysmal asymmetrical facial grimacing was greatly reduced in mice that received Gi DREADD but not mice received vector control (**Figure 4c to f, Supplementary Figure 6a to c**). We also examined relief of pain by conditional place preference (**Figure 4g**), which has been increasingly used as a reliable measurement of pain relief^31^. Results showed that mice received Gi DREADD displayed significant preference in C21-paired chamber, but not saline-paired chamber (**Figure 4h and i**). On the other hand, mice received vector control did not display preference of any chamber, supporting that inhibiting c-Fos expressing neurons in the S1ULp-S1J alleviated spontaneous pain as well as other pain - related behaviors.

**Figure 4.**
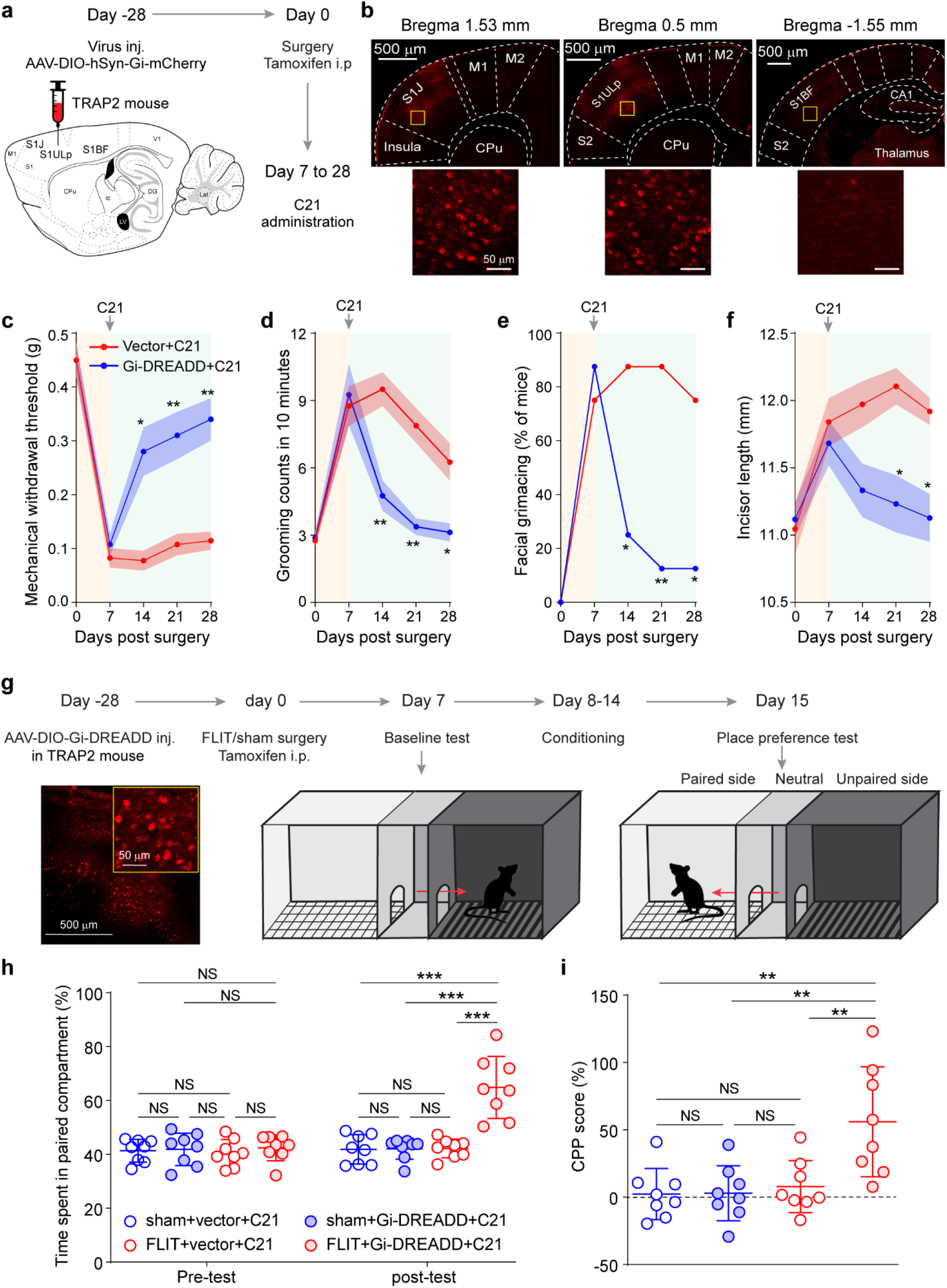
Pain-related c-Fos expressing neurons are critical for pain-like behaviors. **(a-f)** TRAP2 mouse was injected with AAV-DIO-hSyn-Gi-mCherry or AAV-DIO-mCherry vector in S1ULp-S1J at day −28 (*n* = 8 per group). FLIT surgery was performed for all mice at day 0 accompanied by tamoxifen administration. From day 7, C21 was intraperitoneally administrated twice daily. Mice were sacrificed at day 28 for brain slices. **(a)** Diagram of virus injection and flowchart of experiment timeline. **(b)** Upper panels: Representative tangential slices of S1J, S1ULp and S1BF demonstrating mCherry expression was primarily located in S1ULp-S1J. Lower panels represent boxed regions of corresponding upper panels. **(c to f)** Chemogenetic inhibition of c-Fos-induced Gi-expressing neurons leads to attenuated pain-like behavior. Behavioral testing was performed at indicated timepoints. At day 7, behavioral testing was performed prior to C21 administration to obtain pre-treatment baseline. Two-way ANOVA followed by Bonferroni post hoc test was carried out to determine the difference between the two groups. Data are represented as mean ± SEM. *\*P* < 0.05, \*\**P* < 0.01. **(c)** Mechanical withdrawal threshold to von Frey filament. **(d)** Facial grooming counts in 10 minutes. **(e)** Percentage of mice with asymmetrical facial grimaces. **(f)** Incisors length. **(g to i)** Chemogenetic inhibition of c-Fos-induced Gi-expressing neurons leads to conditioned place preference. TRAP2 mice were injected with either AAV-DIO-Gi-GREADD or vector virus at day −28. FLIT or Sham surgery was performed at day 0 accompanied by tamoxifen administration. After baseline test at day 7, C21 was intraperitoneally administrated twice daily, followed by conditioning then place preference test at day 15. Two-way ANOVA followed by Bonferroni post hoc test was carried out to determine the difference among groups. Data are represented as mean ± SEM. \*\**P* < 0.01, \*\*\**P* < 0.001, ns: non-significant. **(g)** Flowchart of conditioned preference place (CPP) experiment. Left lower panel: representative images of c-Fos-induced mCherry-expressing neurons in S1ULp-S1J. **(h)** Percentage of time spent in C21-paired compartment. **(i)** CPP score.

Although c-Fos expression was primarily induced in the S1ULp-S1J region, there was noticeable expression in the S1BF. To examine if those c-Fos^+^ neurons in the S1BF were also implicated in spontaneous orofacial pain, TRAP2 mice were microinjected with Gi DREADD or vector control in S1BF followed by FLIT surgery and C21 injection (**Supplementary Figure 6d and e**). Results showed that twice daily C21 injection in the Gi DREADD group did not significantly alleviate spontaneous pain, as determined by spontaneous facial grimacing (**Supplementary Figure 6f to l**).

### Local GABAergic neuron hypoactivity promotes S1 neural synchronization

Cortical GABAergic interneurons are critical components of cortical local circuits and exert indispensable roles in modulating the timing, extent, and propagation of excitatory neuronal activities^32^,^33^. We examined GABAergic neurons in the FLIT model by taking advantage of a pan-interneuron targeting AAV-hDlx-GCaMP6f for calcium imaging^34^. This tool was able to target about 85% of all Gad67+ interneurons as reported (**Supplementary Figure 7a and b**). Intravital two - photon calcium imaging was performed in Layer 2/3 neurons of S1ULp - S1J in awake mice (**Figure 5a**). Results showed that for the GABAergic interneurons, FLIT group displayed significantly lower total integrated calcium activities than control group (**Figure 5b and c**), suggesting hypoactivity of these local GABAergic interneurons. RNA Seq was performed in S1ULp - S1J in the FLIT and Sham groups, results showed a dramatic decrease in GABA - related genes (**Figure 5d, Supplementary Figure 7c to e**), suggesting GABAergic interneuron hypoactivity was implicated in excitatory neuron synchronization and pain.

**Figure 5.**
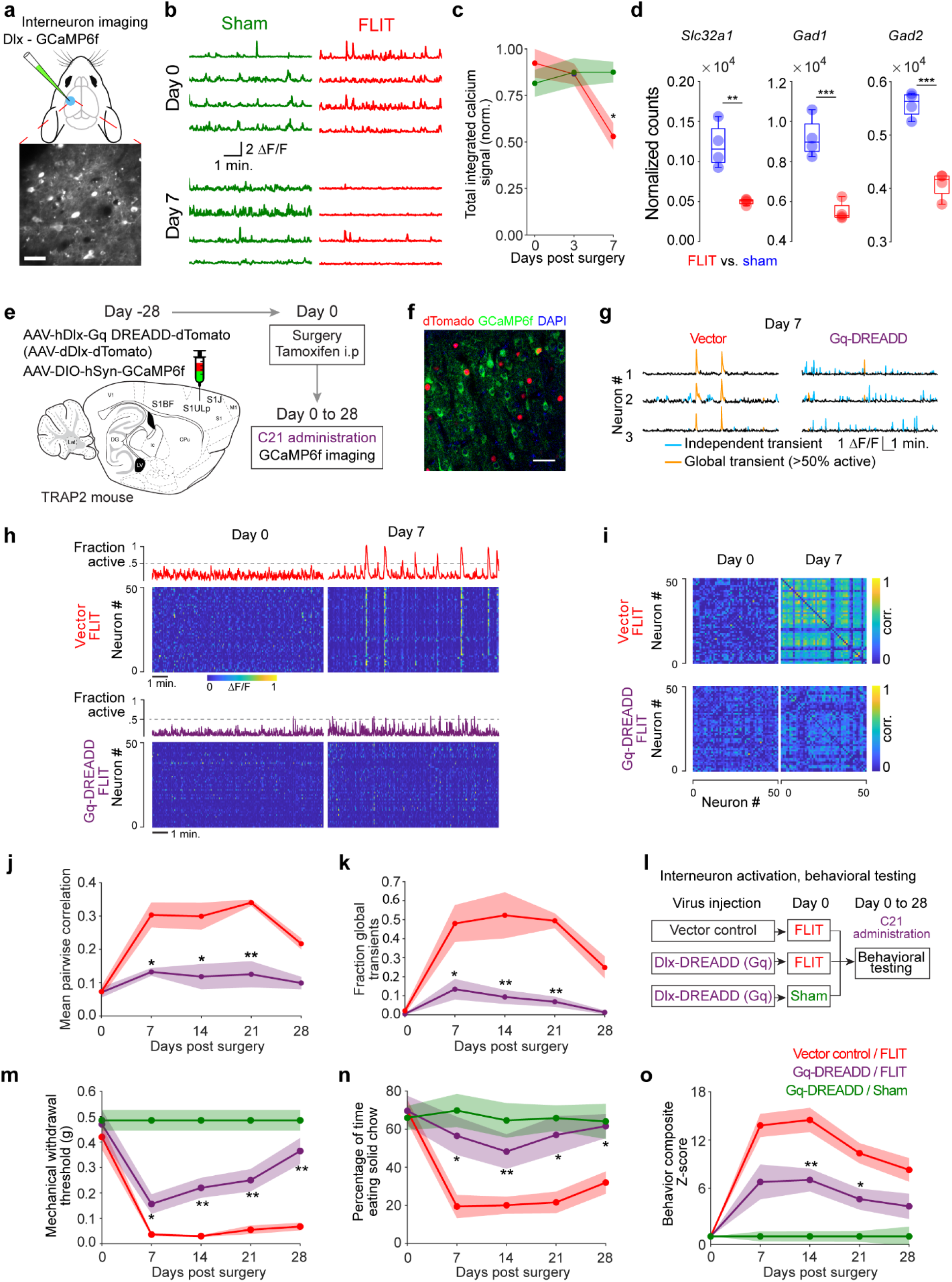
Dampening S1 synchronization through interneuron activation alleviates pain-like behavior. **(a to c)** Interneuron calcium imaging. S1ULp-S1J region was injected with AAV-Dlx-GCaMP6f at day −28, followed by FLIT or Sham surgery at day 0. Calcium imaging was performed at indicated timepoints (*n* = 4 per group). **(a)** Diagram of S1 injection. Lower panel is a representative GCaMP6f expression from an imaging field. **(b)** Representative single neuron calcium dynamics tracing at days 0 and 7 for FLIT and Sham groups. **(c)** Total integrated calcium signals of imaged interneurons. Two-way ANOVA: *P* < 0.05, post-hoc Tukey-Kramer test shows significant difference between FLIT and Sham group, \**P*<0.05. **(d)** Comparison of GABAergic interneuron-related gene transcripts. **(e to k)** Chemogenetic activation of interneuron dampens synchronization of c-Fos-induced pain-related neurons. TRAP2 mice were injected with AAV-Dlx-Gq DREADD-dTomato or AAV-Dlx-dTomato (vector) mixed with AAV-DIO-hSyn-GCaMP6f at day-28 (sham *n* = 6, FLIT *n* = 5). At day 0, all mice underwent FLIT surgery accompanied by tamoxifen administration. C21 was administered twice daily for chemogenetic activation. **(e)** Diagram and flowchart of experimental design. **(f)** Representative brain slice expressing dTomado and GCaMP6f within S1ULp-S1J. **(g to i)** Chemogenetic activation of interneurons decreases synchronization of c-Fos induced pain-related neurons. **(g)** Representative calcium dynamic tracing at day 7 after FLIT surgery in vector and Gq-DREADD group. **(h)** Representative heatmaps and fraction of active neuron plots. **(i)** Representative correlation matrix plots. **(j)** Mean pairwise correlation at different timepoints after FLIT surgery. **(k)** Fraction of global transients among total transient at different timepoints after FLIT surgery. Two-way ANOVA test indicates significant difference between the groups. Post-hoc Tukey-Kramer test was carried out to determine the *P* value of vector / FLIT *vs*. Gq-DREADD / FLIT, * *P* < 0.05, ** *P* < 0.01. (**l to o)** Chemogenetic activation of interneurons alleviates pain-like behavior (*n* = 7 Sham, *n* = 8 for other groups, mean ± SEM). **(l)** Flowchart of experimental design. **(m)** Mechanical withdrawal threshold to von Frey filament. **(n)** Solid food preference. (**o)** Composite Z scores of behaviors (mechanical withdrawal; grooming; body weight; incisor length; wood weight changes; and solid food preference) were computed for all groups. For (m-o), two-way ANOVA test indicates a significant difference among the groups. Post-hoc Bonferroni test was carried out to determine the *P* value of Vector / FLIT *vs*. Gq-DREADD/FLIT, \**P* < 0.05, \*\**P* < 0.01.

To directly examine the functional significance of GABAergic interneurons in synchronized neural dynamics and pain, AAV-hDlx-Gq DREADD-dTomato (Gq DREADD) was used to activate GABAergic interneurons, with AAV-dDlx-dTomato as vector control. For imaging studies, S1ULp-S1J region of TRAP2 mice were microinjected with a mixture of AAV-hDlx-Gq DREADD-dTomato and AAV-DIO-hSyn-GCaMP6f or a mixture of AAV-dDlx-dTomato (vector) and AAV-DIO-hSyn-GCaMP6f, followed by FLIT surgery in 4 weeks (**Figure 5e and f**). Tamoxifen was administered to induce Cre expression, and C21 was injected twice daily to activate Gq DREADD. When two-photon calcium imaging was performed in awake mice, mice received vector control displayed robust synchronization as observed in Figure 3. However, in mice received Gq DREADD, the synchronization was nearly abrogated (**Figure 5g to k**). Thus, activating S1ULp-S1J GABAergic neurons abrogated neuronal synchronization in S1.

To assess if abrogating neuronal synchronization would also alleviate pain-like behaviors, mice with microinjected with AAV-hDlx-Gq DREADD-dTomato (Gq DREADD) or AAV-dDlx-dTomato vector control (**Figure 5i, Supplementary Figure 8a**). Following FLIT surgery, C21 was used to activate DREADD. Remarkably, mice received Gq DREADD exhibited attenuated pain-related behaviors, including spontaneous pain-like behavior, whereas mice received vector control displayed spontaneous paroxysmal asymmetrical facial grimaces. Moreover, global assessment of pain was also carried out using composite pain scores, and results showed that Gq DREADD group had significantly lower pain scores (**Figure 5m to o; Supplementary Figure 8 b to d**). As such, activating S1ULp-S1J GABAergic neurons alleviated pain-like behavior, including spontaneous pain.

### Attenuation of S1 neural dynamics with clinically effective pain treatment

We next asked if clinically effective pain treatment would attenuate S1 synchrony. Carbamazepine, an anti-seizure medication, is currently used clinically as first-line treatment for TN^35^. We therefore applied it to FLIT model mice (**Figure 6a**). S1 synchronization was significantly decreased by carbamazepine (**Figure 6b and c**). Carbamazepine also dampened mechanical allodynia, reduced facial grooming frequency, modestly reduced facial grimaces albeit not statistically significant (**Figure 6d to f**), consistent with clinical reports of partial pain resolution with carbamazepine. In contrast, ketorolac, a non-steroidal anti-inflammatory medication with no clinical efficacy against TN^36^, did not significantly alter neuronal activity or alleviate pain-like behavior (**Supplementary Figure 9a to i**).

**Figure 6.**
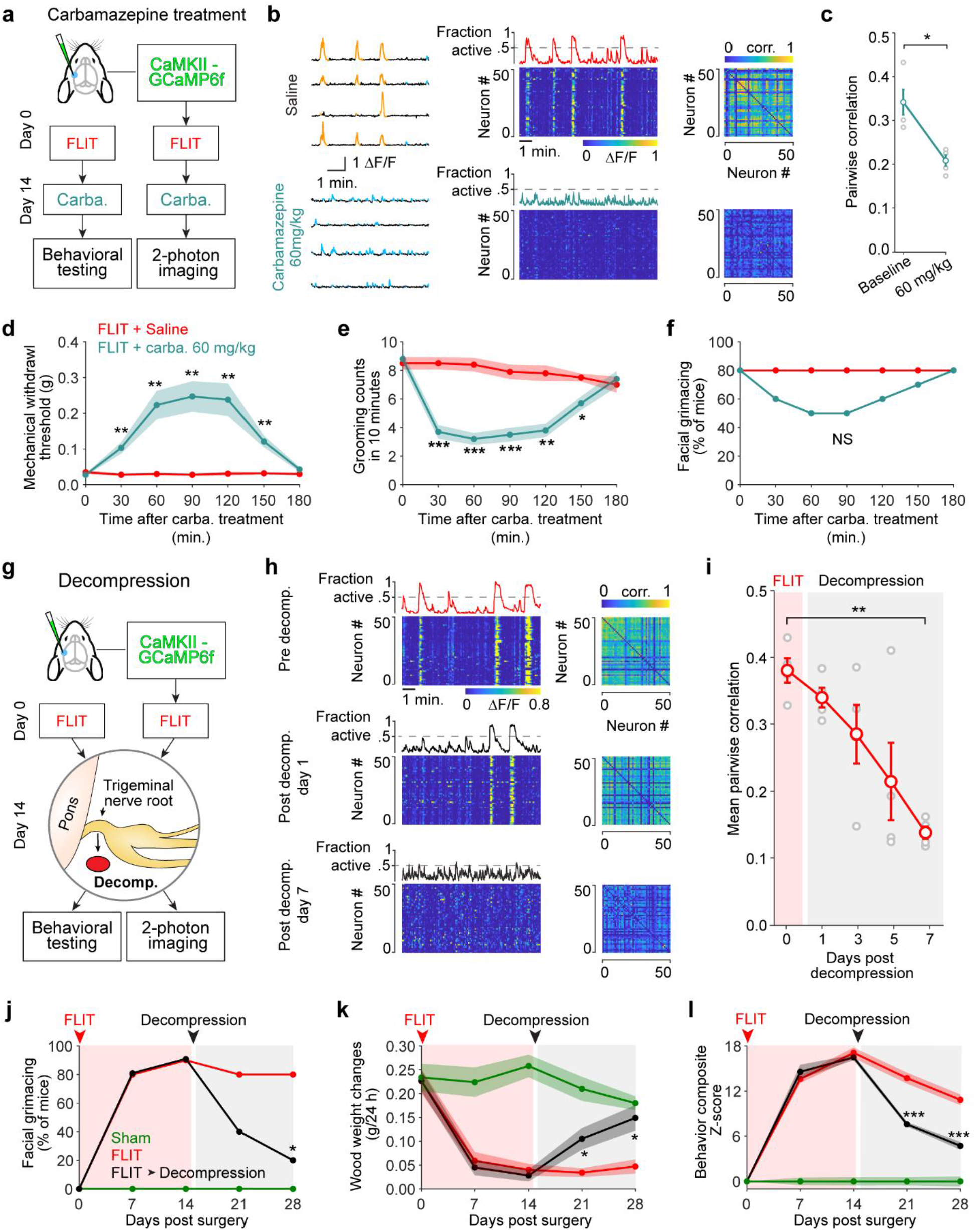
Clinically effective treatments alleviate S1 synchronization and pain-like behavior. **(a)** Diagram depicting carbamazepine experiment. Excitatory neuron imaging and behavioral testing was carried out on separate groups of animals. **(b and c)** Two-photon calcium imaging 14 days after FLIT surgery, animals (*n* = 4) received normal saline followed by carbamazepine 60 mg/kg with a 12-hour interval. **(b)** Left: Sample calcium transient traces of neurons after carbamazepine or saline injection. Middle: heatmap of neuronal activity. Right: correlation co-efficient matrices of neurons shown in the left. Images were acquired from the same field of view but not necessarily the same neurons. **(c)** Pairwise correlation for each treatment for all four animals were shown (gray circle). Mean ± SEM, Paired *t*-test: \**P* < 0.05. **(d to f)** Fourteen days after FLIT surgery, animals received normal saline followed by carbamazepine with a 12-hour interval (*n* = 10). **(d)** Mechanical withdrawal threshold to von Frey filaments (mean ± SEM). Two-way ANOVA test indicates significant difference between the two groups, post-hoc Bonferroni test indicates different at indicated time points saline *vs*. carbamazepine, \*\**P* < 0.01. **(e)** Facial grooming counts (in 10 minutes) after saline or carbamazepine treatments. Two-way ANOVA test indicates significant difference between the two groups, post-hoc Bonferroni test was carried out to determine the *P* value of saline *vs*. carbamazepine, \**P* < 0.05, \*\**P* < 0.01, \*\*\**P* < 0.001. **(f)** Percentage of mice with asymmetrical facial grimaces. No statistical significance was present between the groups (Fisher’s exact test). **(g)** Flowchart of experiment of S1 calcium imaging (*n* = 4) and behavioral tests (*n* = 10 per group) for decompression (removal of Surgifoam) of trigeminal nerve root in FLIT mice. **(h)** Representative neuronal activity heatmaps and representative correlation co-efficient matrices from the same animal before and after decompression. **(i)** Pairwise correlation co-efficient before and after decompression (individual animals in gray, mean ± SEM in red). One-way ANOVA \*\**P* < 0.01, Tukey-Kramer post-hoc comparison shows a significant difference between day 0 and day 7. **(j to l)** Behavioral tests of mice underwent decompression at 14 days post FLIT surgery. **(j)** Asymmetrical facial grimaces. \**P* < 0.05 by Fisher’s exact test. **(k)** Wood chewing assay. Two-way ANOVA test indicates significant difference among the groups, post-hoc Bonferroni test was carried out to determine the *P* value of FLIT *vs*. FLIT + decompression, \**P* < 0.05. **(l)** Composite Z-score of behaviors were computed for all groups. Two-way ANOVA test indicates significant difference was present among the three groups. Post-hoc Bonferroni test was carried out to determine the *P* value of FLIT *vs*. FLIT + decompression, \*\*\**P* < 0.001.

We tested if trigeminal nerve root decompression surgery (**Figure 6g**), a definitive treatment for TN patients with vascular compression^37^, altered cortical S1 activity dynamics. Population dynamics in S1 showed strongly attenuated synchrony after decompression. By day 7 after decompression, S1 neuronal activity returned to levels similar to pre-FLIT baseline (**Figure 6h and i**). The decompressed FLIT mice exhibited attenuated mechanical allodynia: mechanical withdrawal threshold returned to pre-FLIT baselines within 14 days post decompression (**Supplementary Figure 10a**). Consistent with this, asymmetrical facial grimaces, decreased wood gnawing (**Figure 6j and k**), facial grooming, incisors overgrowth and body weight were also reversed (**Supplementary Figure 10b to d**). Composite Z-score derived from pain-related behaviors revealed that decompression reversed the behavioral phenotypes within 14 days (**Figure 6l**). We also performed a recompression experiment in decompressed mice that had resolved cortical synchrony (**Supplementary Figure 10e and f**) and observed reoccurrence of synchronized S1 activities following recompression, confirming the causal link between induction of pain and synchronized S1 activity.

## Discussion

We showed a dramatic increase in synchronicity of S1 pyramidal neurons as a result of neuropathic pain (Figure 2 and Supplementary Figure 5). This hypersynchrony is not a mere by-product of pain, rather, it is critical for pain-like behavior, including spontaneous pain. The interrogation of cortical mechanisms responsible for spontaneous pain was greatly facilitated by our novel, clinically relevant model of TN. We developed this model after reasoning that cortical neuronal dynamics could be well captured in a model of orofacial pain with robust spontaneous pain-like behavior. The orofacial area occupies a disproportionally large region on the S1 homunculus compared to other body parts^20, 21, 38^, enabling the investigation of cortical neural dynamics of pain.

Pathological insults to the nervous system can induce characteristic hypersynchronous states, as reported in Alzheimer’s disease, Parkinson’s disease, and epileptic seizures^39–42^. A mechanistic understanding of aberrant neural synchronization in these pathological conditions is currently lacking. We showed that S1 interneuron hypoactivity is key for excitatory neuron hypersynchrony. Selectively activating interneurons in the affected areas reversed hypersynchrony of excitatory neurons and alleviated pain-like behavior, enabling us to gain insights into the mechanisms underlying the observed pathological activity in pain. Such S1 interneuron hypoactivity has been recently shown to mediate sensory hypersensitivity in a mouse model of autism spectrum disorder^43^ and may indeed underly a range of other pathologies of the brain.

Cortical hypersynchrony observed in neuropathic pain was functionally important for pain-like behavior. Reversing the S1 hypersynchrony through local interneuron activation ameliorated pain-like behavior (Figure 5), indicating that targeting aberrant pain-induced S1 neural dynamics could relieve pain. Interestingly, anti-seizure medication, carbamazepine, which is used as first-line medication for TN, but not ketorolac, a non-steroidal anti-inflammatory drug, could alleviate S1 hypersynchrony. Consistent with this, trigeminal nerve root decompression, a definitive treatment of TN for a subset of patients with surgically-amenable nerve root compression, also attenuated S1 hypersynchrony (Figure 6).

Elevated synchrony of neurons is not necessarily a sign of pathology. Neuronal synchronization in the visual cortex has been shown to establish relations in different parts of the visual field coding for global features of stimuli such as continuity, similarity of orientation and coherency of motion^44^. The olfactory system also demonstrates transiently synchronized neuronal activities in odor-evoked dynamic ensembles. This temporal synchronization is linked to combinatorial representations of time and space during an odor response^45, 46^. More recently, synchronization of the cortical layer 5 pyramidal neurons has been linked to loss of consciousness during general anesthesia^47^. It is therefore important to delineate pathological synchrony from otherwise healthy cortical function. The pain model we developed provides a novel method to reversibly induce pathological hypersynchrony. Most existing pain models derive from original insults that could not be reversed in a temporally controlled fashion. As shown (Figure 6 and Supplementary Figure 10), trigeminal nerve root decompression alleviated cortical hypersynchrony and nerve re-compression re-introduced the hypersynchrony. As such, decompression and re-compression of the FLIT model change cortical dynamics in a ‘predictable’ fashion. Our approach thus establishes a robust and reproducible method for investigating how pain can alter cortical microcircuitry linked to intractable, persistent maladaptive behavior.

Recent research has started to investigate the cortical representation of pain. Formalin-induced pain has been found to be associated with neuronal oscillation in S1^48^. In a mouse spared nerve injury model of neuropathic pain, heightened S1 neuronal activity has been shown to mediate pain perception^19^. However, in these studies, whether or not cortical neurons exhibit synchrony is unclear. Synchronized cluster firing has been reported in anesthetized condition in the dorsal root ganglion primary sensory neurons^49^. Our results indicate that in awake state with no experimental stimuli, localized S1 synchronization is a key cortical pattern mediating pain-like behavior. Connecting these observations yields a new hypothesis that peripheral and central mechanisms orchestrate the neurological manifestations of spontaneous pain.

Using immediate early genes, particularly c-Fos, to map neural activity associated with biological and behavioral perturbations has led to many exciting discoveries^28^, including the discovery of anesthesia-related analgesic effects mediated by a group of neurons in the amygdala^50^. We leveraged c-Fos immunostaining to map pain-related cortical activity in S1ULp - S1J in mice that underwent induction of neuropathic pain. Notably, these mice did not receive any experimentally imposed stimulation to evoke pain responses. Therefore, c-Fos expression attributed to internal processing of pain could be derived by comparing animals that underwent FLIT and Sham surgeries. Consistent with c-Fos expression, neural activity was significantly increased in S1ULp-S1J neurons (Figure 2 and Supplementary 4). Using TRAP2 mice^28^, we interrogated the function of spontaneous pain-related c-Fos expressing neurons and found these neurons were indispensable for mediating spontaneous pain-like behavior. Combining behavioral and imaging approaches, we reported a regional difference between S1ULp-S1J and S1BF in mediating spontaneous pain (Figure 4 and Supplementary Figure 6). S1 BF has been long recognized as a critical region decoding whisker stimulation^51^, our results indicate that cortical processing of spontaneous pain in the orofacial areas involves different mechanisms than vibrissal sensory processing by S1BF. Recently, the dysgranular region was implicated in S1 processing of pain^52^. In our orofacial spontaneous pain model, we did not observe significant c-Fos expression in this region, which might be related to different types of pain models as well as the presence of absence of spontaneous pain^52^.

Taken together, we found distinct states of S1 hypersynchrony dynamics as a key neural substrate mediating spontaneous pain. Our findings provide mechanistic insights into this devastating aspect of human neuropathic pain and open up new avenues for new treatments targeting pathological neural synchrony.

## Materials and Methods

### Animal

All animal use and procedures applied according to protocols approved by the Massachusetts General Hospital Institutional Animal Care and Use Committee (IACUC). Experiments performed were complying with the guidelines established by NIH and the International Association for the Study of Pain. Male *Fos^2A-iCreER^* mice (TRAP2) (Jax 030323), adult male and female C57/BL6 mice (16-24 weeks old) were purchased from the Jackson Laboratory (ME). Mice were housed in temperature-controlled vivarium on a 12 h light/dark cycle (lights on at 07:00 am, lights off at 07:00 pm) with food and water available *ad libitum*. CD1 mice (16-26 weeks old, male) and SD rats (10 to 14 weeks old, male) were also purchased from the Jackson Laboratory (ME) and Charles River Laboratories (MA), respectively. For carbamazepine, oral gavage of carbamazepine (Novartis, packaged by Precision Dose, Inc) at 60 mg/kg was used. For ketorolac (Hospira, IL), oral gavage of 10 mg/kg was used. For tamoxifen (Sigma, Lot# WXBD4583V, St. Louis, MO), intraperitoneal injection of 150 mg/kg dissolved in corn oil was used.

### Foramen lacerum impingement of trigeminal-nerve (FLIT) procedure

Mice were anesthetized with isoflurane inhalation (3% for induction and 1.5 to 2% for maintenance) in oxygen (flow rate: 1 to 2 L/min). Surgery was performed under an Omano surgical microscope (OM2300S-V7) at 7.5-45X magnification. After surgical field preparation, a 1.5 cm midline neck incision was made starting at the rostral end of the sternum using a sterile spring scissor. The superficial tissues such as salivary glands were bluntly dissected and lateralized with a mini retractor. The neck muscles, including sternocleidomastoid muscle, digastric muscle, strap muscles were gently dissected to locate right auditory bulla and auditory capsule on the right side of the mouse head which are the landmarks to locate the foramen lacerum. Electric cauterization was applied to control minor bleeding from capillaries. Care was taken while dissecting the vessels covering both the anterior bulla and the foramen of lacerum. A prepared piece of Surgifoam (Ethicon, Inc., Somerville, USA) at ~1-1.5 mg was gently delivered into the foramen lacerum using curved forceps. The Surgifoam was positioned between trigeminal nerve root and the cochlea bulla. After removing the retractor and replacing the tissues, the skin was closed with 6-0 nylon monofilament (Ethicon, Inc., Somerville, USA) sutures. Mice in sham group underwent the same surgical procedure including neck shaving, skin incision, muscles dissection and foramen lacerum exposure without physical trigeminal nerve root compression. The surgery time ranged from 8-12 minutes per mouse. Tamoxifen was intraperitoneally administrated in Fos-iCre-ERT2 (*Fos^2A-iCreER^* knockin) mice immediately after the FLIT procedure.

### Trigeminal nerve root decompression and recompression

At 14 days post FLIT procedure, mice were anesthetized with isoflurane in oxygen. The same neck incision and neck cervical dissection was performed as described in the FLIT procedure. After locating the foramen lacerum, the Surgifoam was removed with care. The incision was closed as described in the FLIT procedure. Recompression surgery was performed as described in the FLIT procedure.

### Mechanical withdrawal threshold

Mice were individually placed in a custom-made box (6×6×6 cm) with the top, bottom and four walls made of metal mesh and allowed for free movement. After 30 minutes of acclimation, a graded series of von Frey filaments were inserted through the mesh walls from the lateral side and applied to the skin of the vibrissa pad within the trigeminal nerve V2 branch-innervated territory for 1 second at 10 seconds intervals. A brisk withdraw of the head upon stimulation was considered a positive response. Mice were tested 5 times with at least 3 positive responses indicated a positive result. The minimum force necessary to elicit a response was defined as the mechanical withdrawal threshold.

### Observation of face grooming and grimacing

For facial grooming and grimacing test, each mouse was habituated 30 min daily for up to 3 consecutive days in a 10×10×12 cm Plexiglas box equipped with a mirror to record unobstructed views of the orofacial area. The mice behaviors were recorded for 10 minutes without any extra audio or physical disturbance. Grooming was defined as face wash strokes primarily directed to the trigeminal nerve impingement side. Facial grimacing for this study was defined as asymmetrical eyelid contraction that ipsilateral eye (same side as trigeminal nerve compression) opening was smaller than contralateral side as determined by blinded observers. The recorded behaviors were analyzed by an experimenter who was blinded to the procedures condition/group assignment of the mice.

### Food preference

Mice were deprived of food 12 hours prior to the test, with water accessible *ad libitum*. To prepare the soft chow, regular solid chow was soaked in water (pellets: water = 1: ~2 g) for 20 minutes. Regular solid chow and freshly prepared soft chow were placed in plates. Test mouse was videotaped using a camera 40cm above the cage for 10 minutes. Time spent eating solid and soft chow in each video was quantified by experimenters that were blinded to group assignment.

### Wood chewing assay

Balsa wood blocks were customized to 1-inch cubes. Mice were housed in individual cages with food and water supplied *ad libitum*, a wood block was placed in the cage for 24 hours. Weight of blocks before and after placement was recorded.

### Behavioral composite Z-score

For comprehensive assessment of several pain-related parameters, we used a formula Z = [Δ XSurgery-MEAN(Δ X)Sham]/SD(Δ X)Sham^53^. In the formula, Δ XSurgery was the score of mice in the surgery group at different time points minus the score of these mice at day 0 baseline; MEAN(Δ X)Sham was the score of mice in the Sham group at different time points minus the score of these mice at day 0 baseline; and SD(Δ X)Sham was the standard deviation of Δ XSham for any given time point. Specifically, the composite Z-score for the mouse was calculated as the sum of the six values of Z-score (mechanical withdrawal thresholds, grooming count, body weight, wood chewing, incisors length, and food preference were tabulated.) normalized with the SD for that sum in the Sham controls.

### Mouse sexual function

For sexual function, naive female mice were used as mating partners. Male mice were placed in cage with a camera placed 40 cm above to record mating behavior. Mounting time and attempts were obtained from the videos by experimenters who were blinded to study design.

### Testosterone quantification

Urine testosterone quantification was performed according to manufacturer’s recommendation (R&D Systems, CatLog KGE010, MN, USA). Briefly, assay plate was prepared by primary antibodies at room temperature for 1 hour. Urine samples were added to each well and incubated for three hours at room temperature, which was followed by substrate reaction at room temperature for 30 minutes. A microplate reader (450nm) was used to determine the optical density of each well.

### Open field test

Mice were habituated for 30 minutes ahead of the test to allow acclimation of the testing environment. Each mouse was placed in a 40×40-cm wall-enclosed box while concurrently activating SMART video tracking system (SMART software vision 3.0, Panlab now Harvard Apparatus), locomotion activity was recorded for 10 minutes with minimal external stimuli. Behavior parameters, including percentage of time spending in center zone and latency of first entry to center zone, were automatically tabulated by the software and analyzed by experimenter who is blinded to the study design.

### Conditioned Place Preference (CPP)

CPP testing was performed as previously described^54^. Briefly, CPP was performed in a three-chamber apparatus (Med-Associates, Inc, St Albans City, VT) containing a white and a black compartment (20.3 x 15.9 x 21.3 cm) with distinct patterns on the floors, separated by a central grey neutral area. Male *Fos^2A-iCreER^* mice were injected with either AAV8-hSyn-DIO-hM4D(Gi)-mCherry (Addgene 44362) or vector virus AAV8-hSyn-DIO-mCherry (Addgene 50459) at day −28. At day 0, these animals received FLIT or sham procedure, immediately followed by single dose of tamoxifen (150 mg/kg) intraperitoneal injection. At day 7 post FLIT surgery, animals were screened using a pre-conditioning test. During the pre-conditioning test, mice were allowed 10 min free access to all compartments. Mice spent less than 75% of their time in any one compartment were included in the study. Conditioning phase started at day 8 after FLIT surgery. During the conditioning phase, mice were confined to one compartment for 45 min after an injection of C21 (1 mg/kg, intraperitoneal), or to the other compartment after a saline injection with 6 hours of interval. After 7 consecutive days of conditioning, mice were retested. Percent of time spent in the paired compartment was calculated for each mouse as T2/(T1+T2) *100, where T1 and T2 represent the time spent in unpaired and paired compartments respectively. And CPP score was calculated for each mouse as ((W2-W1)/W1) *100, where W2 stands for the percent time spent in the C21 paired compartment during the final test and W1 stands for the percent time spent in the same compartment during the initial test.

### Botulinum toxin-A injection

Mice were briefly anesthetized under isoflurane anesthesia. To be consistent with the FLIT procedure on the right side, masseter muscles on the right side were injected with Botulinum toxin A (Merz Pharmaceutical, NC) at 0.4U in 50 μL. H&E staining atrophy was diagnosed by a pathologist who was blinded to group assignment, using muscle fiber diameter and nucleus position as diagnostic criteria.

### Craniotomy and Virus injection

Cranial windows were implanted on the contralateral side to the FLIT or Sham procedure. Mice were anesthetized with isoflurane (3% for induction and 1.5% for maintenance in oxygen). The eyes were moistened with eye lubricant. To minimize post-op pain, ketorolac tromethamine (Althenex, Schaumburg, IL USA) was administrated (5 mg/kg) intraperitoneally every 24 hours for 3 consecutive days. The fur on the top of the head was shaved between outer canthus and concha, then the mouse was positioned in a stereotactic frame with a head holder. The skin was prepared with Povidone-Iodine solution (Aplicare, INC., Neriden, CT USA) followed by 70% alcohol swab (BD, Franklin Lakes, USA). After Lidocaine (0.2 ml, 1%) infiltration, a skin flap overlying the dorsal skull was removed using micro scissors. Connective tissues and periosteum of the parietal skull was thoroughly cleaned. A 3×3 mm piece of bone was removed to reveal the left anterolateral cortex including S1BF, S1ULp and S1J as determined by stereotactic coordinates following Chen *et al*.^43^ and the dura was kept moist with sterile saline.

For GCaMP6f expression in pyramidal neurons of the targeted cortex, adeno-associated virus AAV8 carrying CaMKII-GCaMP6f (pENN.AAV.CamKII.GCaMP6f.WPRE.SV40. Addgene. #100834. 1×10^12^ genome copies per ml) was injected with Nanoject-III (Drummond Scientific Company, model #3-000-207) at a depth of 200 μm beneath pia surface and virus was slowly injected at 4-5 sites ~1 to 4 mm lateral to midline of skull, and bregma ~1 to −2 mm in WT mice. Cre-dependent GCaMP6f virus (AAV.Syn.Flex.GGaMP6f.WPRE.SV40, Addgene #100833, 1×10^12^ genome copies per ml) was selectively injected within S1J (jaw) and S1ULp (upper lip) cortex coordinated at 0.5 − 1.5 mm to bregma, ~2.8 − 3.5 mm lateral to midline in Fos^2A-iCre^/TRAP2 mice; for S1BF cortex injection, Cre-dependent GCaMP6f virus was injected at bregma −0.5 to −1.8 mm, and 3 − 3.5 mm lateral to midline in Fos^2A-iCre^/TRAP2 mice. For expression of GCaMP6f in GABAergic neurons, GCaMP6f in AAV8 under the Dlx5/6 promoter adeno-associated virus (pAAV-mDlx-GCaMP6f-Fishell-2 (plasmid #83899)) was injected in the same area of S1ULp (upper lip) and S1J (jaw) of WT mice.

For expression of Designer Receptors Exclusively Activated by Designer Drugs (DREADDs) in GABAergic or Glutamatergic neurons, pAAV9-hDlx-GqDREADD-dTomato (Addgene plasmid #83897) or AAV8-hSyn-DIO-GiDREADD-mCherry (Addgene plasmid #44362) were injected to S1ULp and S1J at ~ 2.8 − 3.5 mm lateral to midline, bregma 1.5 to − 0.5 mm. For S1BF injection, AAV8-hSyn-DIO-GiDREADD-mCherry was injected at bregma −0.5 to −1.8 mm, and 3 − 3.5 mm lateral to midline in TRAP2 mouse, Control mice underwent the same procedure was injected the same volume of pAAV9-hDlx-dTomato or AAV-hSyn-DIO-mCherry (vector control) into S1ULp and S1J, respectively.

For cranial window implantation, two circular pre-sanitized glass coverslips sized with 3 mm and 5 mm (Warner Instruments, #64-0700 & 64-0720) in diameter individually were conjoined with optical adhesive (Norland Products, INC. Cranbury, NJ USA. #417). The 3 mm coverslip was laid over the pia surface within the craniotomy and the surrounding skull was covered by the 5 mm coverslip. A custom-designed head plate was adhered over the glass window with adhesive luting cement (C&B-Metabond, #171032. Parkell. Edgewood, NY USA).

For widefield imaging of calcium dynamics, AAV-CaMKII-GCaMP6f was injected across a large area of cortex covering S1ULp, S1J, S1 barrel cortex, S1 forelimb, S1 hindlimb, M1, M2, S2, etc. (midline to 3 mm at bregma ~1.5 mm and to ~4 mm at bregma −2 mm). Four weeks were allowed for virus expression. For the wide-field brain imaging window, a trapezoidal craniotomy from midline to 3 mm at bregma ~2 mm and to ~4.85 mm at bregma −3.5 mm respectively lateral to midline was made, brain cortex was covered by a customized coverslip glass and a metal headplate was attached using adhesive luting cement.

### Two-photon imaging

Prior to imaging session, mice were taken to the two-photon microscope room and placed onto microscope stage using a head fixation device for 30 min daily, for 5 − 7 days. In vivo two-photon imaging was performed with a two-photon system (Ultima; Bruker) equipped with a Mai tai laser (Spectra Physics, KMC 100). The laser was tuned to 910 nm and the average laser power through the transcranial window was ~20-30 mW for both excitatory neuron and interneuron imaging acquisition using a 20X, 1.0 NA water-immersion objective (Olympus, Japan). A regular 4x objective (Olympus, Japan) was used for widefield view of S1 cortex imaging. All images except for pilocarpine experiment were acquired at frame rate of 6-12 Hz for about 15 minutes using Prairie View Software in awake status without anesthesia. Three Hz frame rate was used for widefield S1 cortex imaging using a 4x objective. For mice that received pilocarpine, images were acquired for about 3 minutes instead of 15 minutes, to minimize restraining animals during epilepsy. Of note, for any given animal, the field of view was kept constant, to obtain images on the same group of neurons longitudinally.

### Calcium imaging data analysis

Imaging data was corrected for motion between frames using the NoRMCorre software package^55^. Neuron selection was carried out subsequently using custom written software in Matlab (Mathworks). Calcium fluorescence signals of each individual neuron were extracted from the corrected video files. The signal for each neuron was corrected for background fluorescence changes by subtracting the fluorescence changes from the immediate surrounding. Each neuron’s activity time course was then quantified using the formula Δ*F* = (*F* – *F0*) / *F0* where F is the fluorescence signal at a given frame and F0 was calculated from a sliding window of +/-30 seconds around the frame. Finally, baseline correction was carried out by fitting a linear function (Matlab function robustfit) to the lowpass filtered (cutoff: 0.3 Hz) signal. A deconvolution algorithm (Fast online deconvolution of calcium imaging data) was applied to detect transients^56^. The start and end of transients were detected when the model was above 0.1.

Global events were detected when the fraction of simultaneously active neurons exceeded 50% of all neurons in that recording in a given frame. Active neuron refers to a neuron exhibiting a transient that was detected at that frame. Subsequently, a transient was categorized as ‘global’ if any part of it overlapped with a global event. Otherwise a transient was categorized as independent. The fraction of global transients was calculated by dividing the number of global transients by the total number of transients of a given neuron. Only neurons with at least 5 detected transients in a given recording were included in this analysis. The fraction of global transients per neuron was calculated in the same way as previously with normalization for each recording. A second order polynomial function was used to fit to the neurons of each animal.

Pairwise correlation analysis was carried out by calculating Spearman rank correlation coefficient for each pair of neurons in a recording. For each correlation matrix, the autocorrelation (i.e. the correlation of a neurons with itself) was not included in any analysis and is shown in dark blue in the correlation matrices. Average correlation values are calculated as the mean of all pairwise correlations in a recording.

Network analysis was carried out using principal component analysis (Matlab function pca, using SVD) and retaining only the first 3 principal components. Periods where a global event was detected are highlighted in golden. The distance of the network trajectory from the mean was evaluated by taking the mean value for each coefficient and calculating the Euclidean distance of each timepoint to the mean. This was done for timepoints that belong to a global event and those that did not.

Widefield calcium imaging using 4x lens was analyzed by extracting signals of each pixel in the imaging field, the *ΔF/F* for each frame was calculated for each pixel without drawing region of interest. The heatmap was generated by averaging the *ΔF/F* for each pixel over time.

Total integrated calcium activity was quantified by calculating the mean total area under the curve for all neurons in a recording. AUC was normalized for each animal and the relative change across days was calculated as the mean across animals.

All codes are available upon request.

### c-Fos staining

Mice underwent surgeries (Sham and FLIT) were briefly maintained under isoflurane anesthesia and procedures were performed after lidocaine 1% infiltration of incision sites. After two hours recovery, mice were sacrificed and immediately perfused with ice-cold PBS followed by 4% paraformaldehyde in 0.1 M phosphate buffer (4% PFA). Brain samples were fixed in 4% PFA at 4°C for 24 hours. Coronal brain sections were sliced at 60μm thickness using a Leica vibratome (VT 1000s). Slices from bregma 1.6 mm to −1.8 mm were obtained to cover the anterolateral S1 including S1J, S1ULp and S1BF cortex. Ten slices covering anterolateral S1J (bregma 1.6 to 1 mm), fifteen slices covering S1ULp (bregma 1 to 0 mm), and fifteen slices covering S1BF (bregma −0.8 to −1.8 mm) were used for c-Fos combined VGLUT1, VGLUT2 or GAD67 staining.

For the double staining, slices were washed in PBS 5 min for 3 times, followed by blocking with 6% goat serum and 2% bovine serum albumin (BSA) in PBS with 0.3% Triton X-100 (Blocking solution) at room temperature for 1 h. Floating slices were stained with primary antibody (rabbit anti-c-Fos, CellSignaling, CatLog 2250, 1:500 dilution; anti-VGLUT1, Invitrogen, Lot# WH3344773B, 1:500 dilution; Guinea pig anti-VGLUT2, EMD Millipore Corp., Lot# 3482777, 1:500 dilution; anti-GAD67, EMD Millipore Corp., Lot# 3281595) in blocking solution at 4°C for overnight. After washing with PBS 5 minutes for 3 times, slices were incubated with secondary antibody (goat anti-rabbit Alexa488, Jackson ImmunoResearch, CatLog 111-545-144, 1:2000 dilution; Donkey anti-mouse Cy3, Jackson ImmunoResearch, Lo#125797, 1:2000 dilution). Images were acquired with Nikon A1 confocal microscope equipped with 20x and 4x objectives. The acquired images were analyzed using ImageJ (NIH opensource software). Number of positive cells were counted by an experimenter who was blinded to group assignment. For each group, 4 animals were used. Number of c-Fos, GLUT1 and GAD67 positive cells were counted within the slices covering S1J, S1ULp and S1BF individually.

### DREADD expression and GCaMP staining

For expression of Designer Receptors Exclusively Activated by Designer Drugs (DREADDs) in GABAergic or Glutamatergic neurons, pAAV9-hDlx-GqDREADD-dTomato (Addgene plasmid #83897) or AAV8-hSyn-DIO-GiDREADD-mCherry (Addgene plasmid #44362) were injected to S1ULp and S1J at ~ 2.8 - 3.5 mm lateral to midline, bregma 1.5 to - 0.5 mm. For S1BF injection, AAV8-hSyn-DIO-GiDREADD-mCherry was injected at bregma −0.5 to −1.8 mm, and 3 - 3.5 mm lateral to midline in TRAP2 mouse. To determine the DREADD expression, mice were sacrificed and immediately perfused with ice-cold PBS followed by 4% paraformaldehyde in 0.1 M phosphate buffer (4% PFA). Brain samples were fixed in 4% PFA at 4°C for 24 hours. Coronal brain sections were sliced at 50-70 μm thickness using a Leica vibratome (VT 1000s). dTomato or mCherry expression site was examined by imaging the coronal sections from bregma 1.6 mm to −1.8 mm using a wide-field fluorescence microscope (Olympus, Japan), and images scanned with a standard 10x objective. For GCaMP staining, slices were washed in PBS 5 min for 3 times, followed by blocking with 6% goat serum and 2% bovine serum albumin (BSA) in PBS with 0.3% Triton X-100 (Blocking solution) at room temperature for 1 h. Floating slices were stained with primary antibody (mouse anti-GFP polyclonal, Invitrogen, CatLog A6455, 1:500 dilution) in blocking solution at 4°C for overnight. After washing with PBS 5 minutes for 3 times, slices were incubated with secondary antibody (goat anti-rabbit Alexa488, Jackson ImmunoResearch, CatLog 111-545-144, 1:2000 dilution). DREADD (dTomato) and GCaMP expression was examined under confocal microscope (Nikon) using excitation wavelength of 554nm and 488 nm, respectively.

### Gi and Gq DREADD activation for behavioral studies

Intraperitoneal injection of C21 (Tocris, CatLog 5548, MN) was used, dosing regimen was determined according to experimental design.

## Supporting information

Excessive facial grooming in FLIT model

Paroxysmal asymmetrical facial grimacing in FLIT model

Synchronized S1 neuronal activity in the same field of view of a FLIT mouse

## Acknowledgements

The authors acknowledge MGH IACUC and the animal facility for kind support; Scot Mackeil from MGH Bioengineer Lab for anesthesia equipment validation; MGH Neuroscience Center Drs. Yinghua Jiang, Xiaoying Wang, Tom Qin, and Yi Zheng for surgical microscope; and MGH two-photon facility Drs. Yaseen Abbas, Sava Sakadzic, and Caroline Magnain for equipment maintenance and technical support; Dr. Qiufu Ma at Dana Farber Cancer Institute for critical comments on the manuscript. Source of mouse silhouettes: scidraw.io.

## Author contribution

Conceptualization: W.D; L.F; Q. C., M.H; S.S.; Methodology: W.D.; L.F.; Q.C.; Z.L.; P.J.; G.F.; M.H., S.S.; Investigation: W.D.; L.F.; Q.C.; L.Y.; Z.Y.; K.H.; X.W.; X.Z.; L.C.; S.W.; S.X.; P.H.; W.C.; S.Z.; C.B.; D.D; T.D; C.W; B.W. Formal analysis, data curation: W.D.; L.F.; Q.C.; Z.L.; L.Y.; P.J.; M.H.; S.S.; Writing: Original draft: W.D.; L.F.; Q.C.; M.H.; S.S.. Funding acquisition: S.S..

## Data Availability

The datasets generated and/or analyzed during the current study are available from the corresponding authors upon reasonable request. Two-photon imaging pipeline will be deposited to GitHub.

## Supplementary materials

Materials and methods

Figs. S1 to S10

Movies S1 to S3

---

**Figure. S1.**
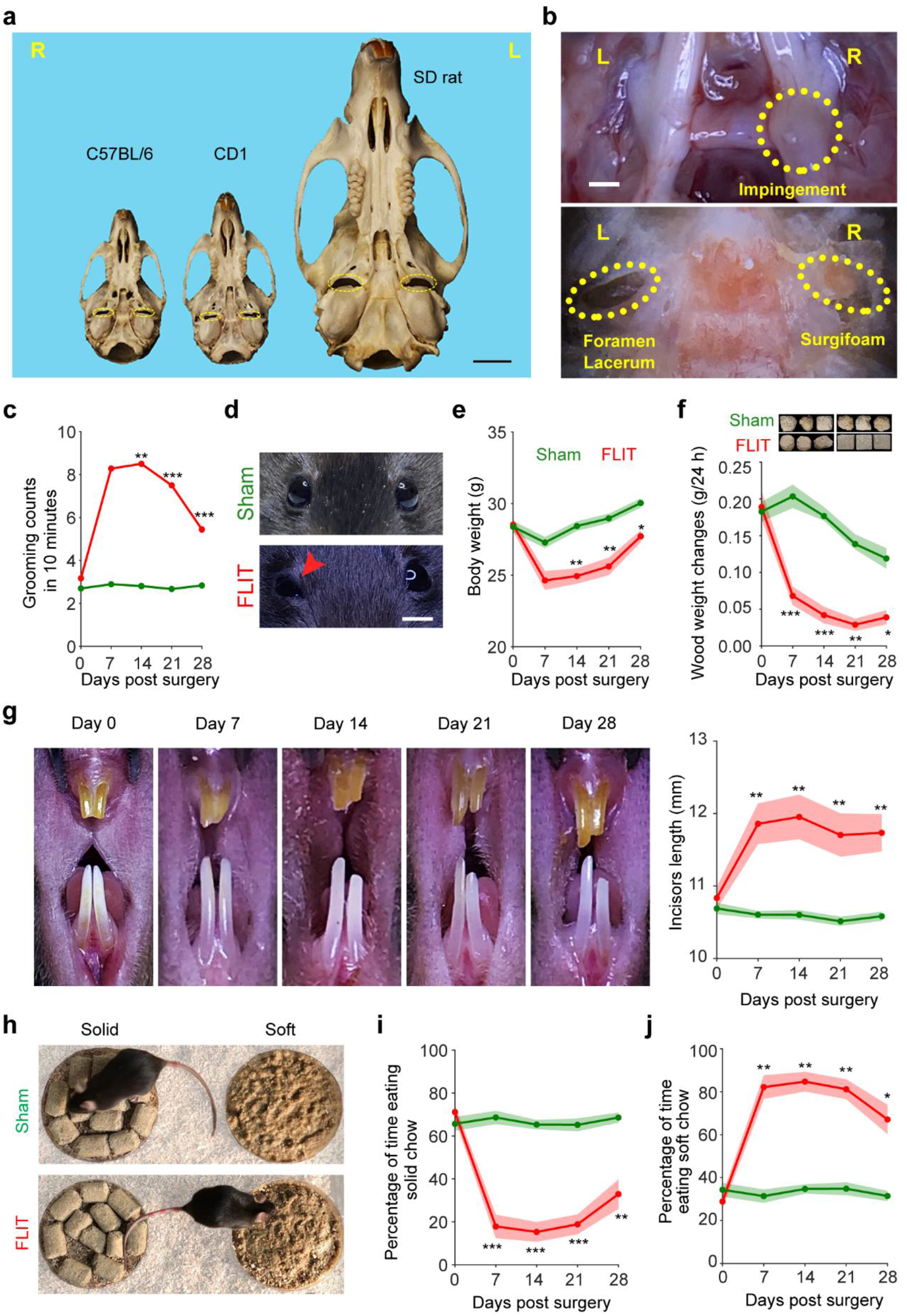
FLIT model. **(a)** Skull base of different species with foramen lacerum labeled in yellow ellipses. Scale bar represents 5 mm. **(b)** A representative picture of trigeminal nerve root impingement from a mouse sacrificed at day 28 post FLIT surgery. Upper panel: top-down view with brain removed to show impingement of trigeminal nerve root on the right side. Lower panel: top-down view with brain removed and trigeminal nerve root lifted to show the Surgiform remained in situ at day 28 post FLIT surgery. Scale bar represents 1 mm. **(c to j)** Behavioral testing for the FLIT model. Mice underwent Sham and FLIT surgery followed by behavioral testing at indicated time points. Equal numbers of male and female mice were used for each group (*n* = 18, Mean ± SEM.). **(c)** Facial grooming counts (in 10 minutes) at indicated time points. Two-way ANOVA indicates significant difference present among the groups, post-hoc Bonferroni test indicates *P* values of FLIT *vs*. other groups, \*\**P* < 0.01; \*\*\**P* < 0.001. **(d**) Representative pictures of mouse eyes among groups. Asymmetric eye grimacing was present only in the FLIT group. Scar bar represents 5 mm. **(e)** Body weight of mice were examined at indicated time points for all groups. Two-way ANOVA indicates significant difference present among the groups, post-hoc Bonferroni test indicates the *P* value of FLIT *vs*. IoN-CCI \**P* < 0.05; \*\**P* < 0.01. **(f)** Top: A representative picture of wood chewing assay at baseline (day 0) and day 7 post surgery. Scale bar represents 1 inch. Bottom: Wood chewing assay to measure wood weight changes for animals that underwent Sham or FLIT surgery. FLIT mice exhibited significant less wood chewing activity compared with sham mice at days 7, 14, 21, and 28. Tw0o-way ANOVA indicates significant difference among the groups, post-hoc Bonferroni test was carried out to determine the *P* value of FLIT *vs*. the other groups, * *P* < 0.05; ** *P* < 0.01; *** *P* < 0.001. **(g)** Left panel: Representative pictures of incisors taken for the same mouse to demonstrate incisors overgrowth in the FLIT model. Scale bar represents 1 mm. Right panel: Quantification of incisors length. Two-way ANOVA indicates significant difference present among the groups, post-hoc Bonferroni test was carried out to determine the *P* value of FLIT *vs*. sham group, \*\**P* < 0.01. **(h)** Pictures of food preference assay. **(i)** Percentage of time eating solid chow. FLIT mice exhibited significantly less solid chow eating compared with sham mice at days 7, 14, 21, and 28. **(j)** Percentage of time eating soft chow. FLIT mice exhibited significantly more solid chow eating compared with sham mice at indicated timepoints. Two-way ANOVA followed by post-hoc Bonferroni test was carried out to determine the *P* value of FLIT *vs*. sham, \**P* < 0.05; \*\**P* < 0.01, \*\*\**P* < 0.001).

**Figure S2.**
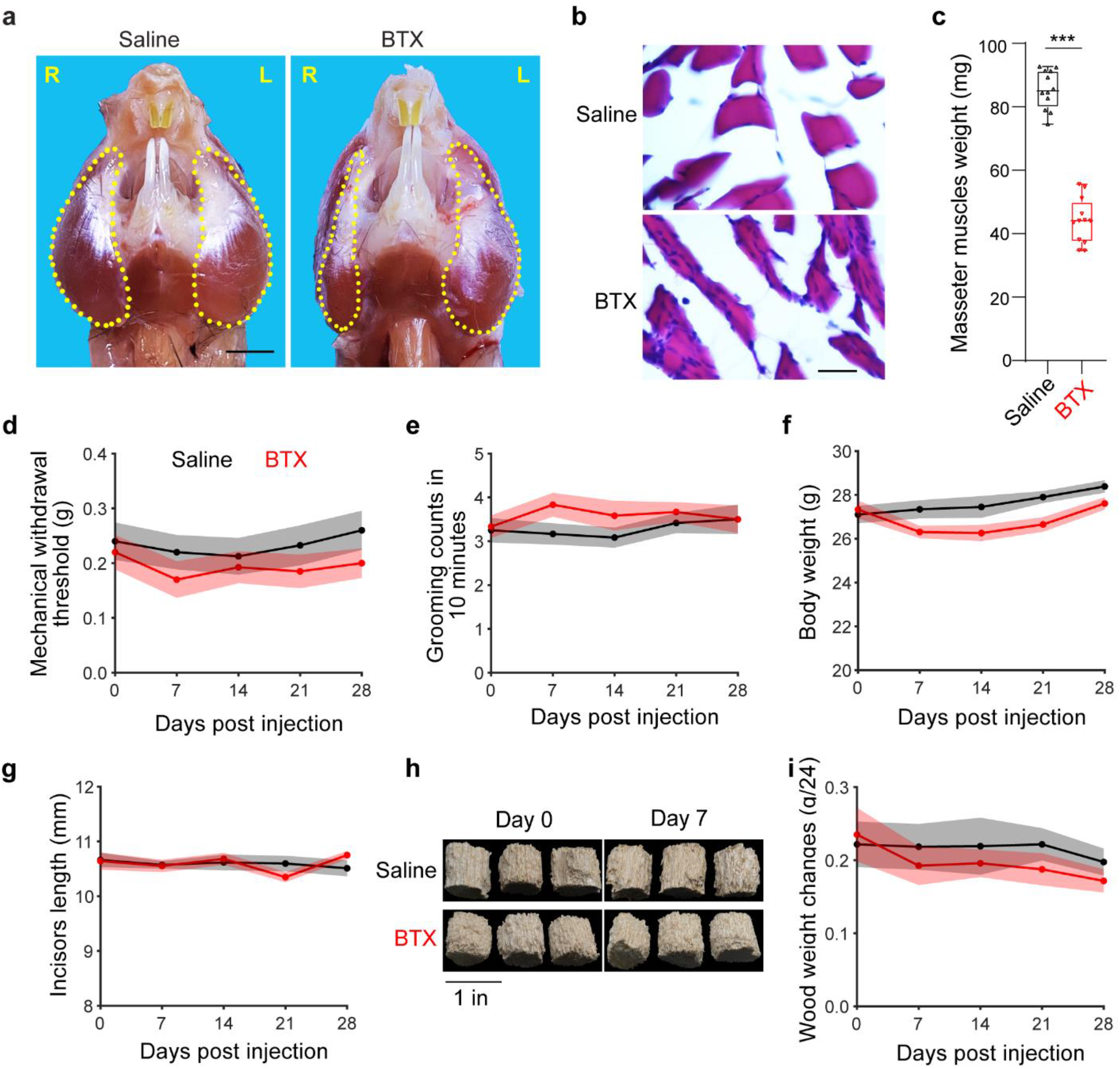
Masseter muscle atrophy does not lead to TN-like behaviors. **(a)** Representative pictures of mouse masseter muscle 28 days post injection with 50 μl saline (left panel) or 50 μl Botulinum toxin-A (0.4U) (right panel). Both saline injection and Botulinum injection were performed on the right side, consistent with the FLIT procedure. *n* = 12 per group. Scale bar represents 5 mm. **(b)** Masseter muscle atrophy revealed by H&E staining. Representative H&E staining of masseter muscle shown (40X). Scale bar represents 100 μm. **(c)** Masseter muscles weight was examined at 28 days after injection. Unpaired *t* test with two-tailed *P* value indicates significant difference present between the two groups (\*\*\**P* < 0.001). **(d-i)** Behavioral testing for the mice received saline or BTX-A injection. *n* = 12 per group. **(d)** Mechanical withdrawal thresholds to von Frey filaments (mean ± SEM). Two-way ANOVA indicates no significant difference between the two groups. **(e)** Facial grooming counts in 10 minutes at indicated time points. Two-way ANOVA indicates no significant difference between the two groups. **(f)** Body weight of mice received saline or BTX-A injection were examined at indicated time points. Two-way ANOVA test indicates no significant difference between the two groups. **(g)** Incisors length of mice were quantified at indicated time points. Two-way ANOVA test indicates no significant difference between the two groups. **(h)** Representative pictures of chewed balsa wood blocks at indicated time points. **(i)** Quantification of balsa wood weight changes. Two-way ANOVA test indicates no significant between the two groups.

**Figure S3.**
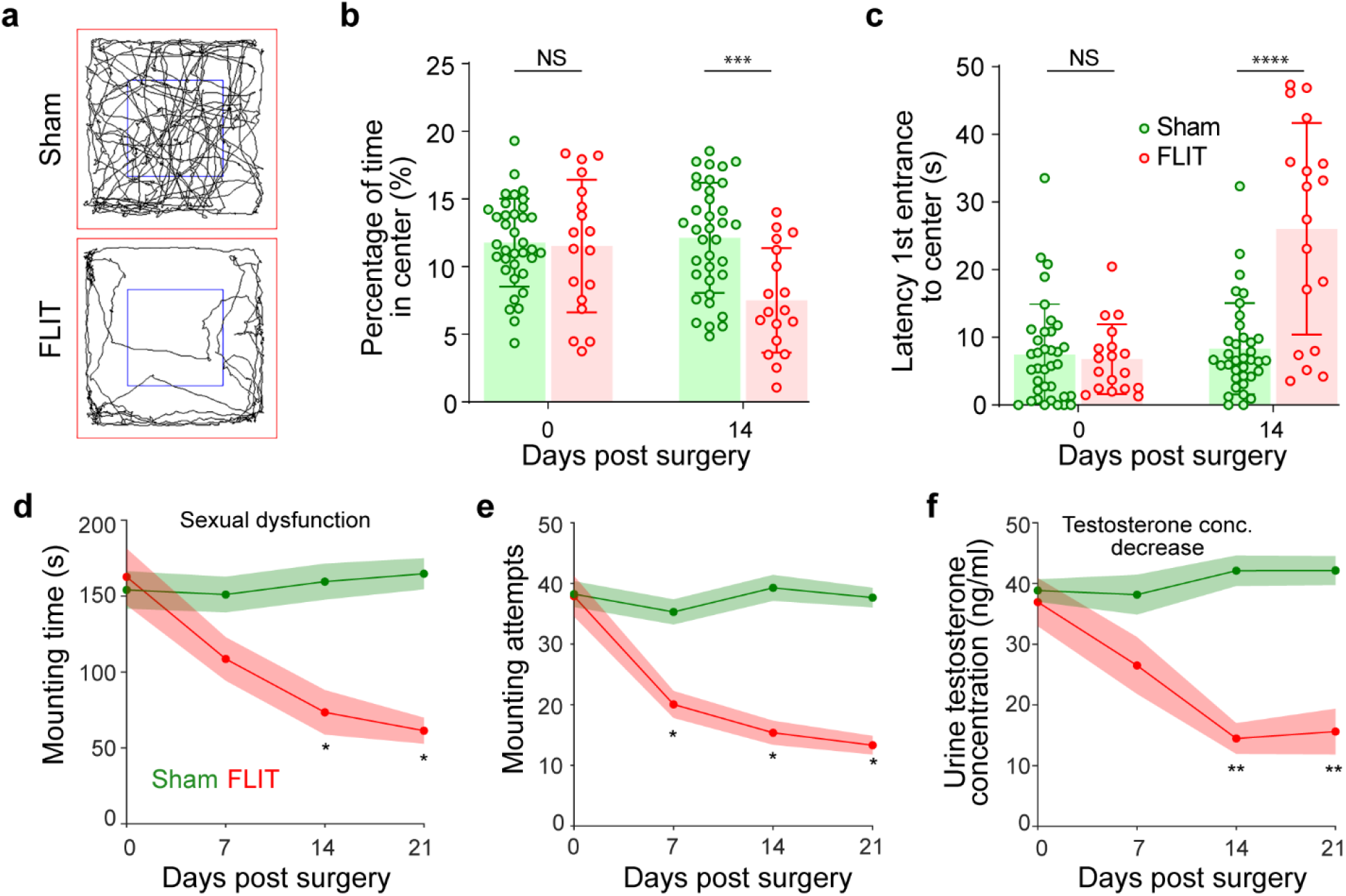
FLIT model develops anxiety-like behaviors and sexual dysfunction. **(a to c)** Anxiety-like behavior was assessed using open-field assay for mice 14 days post surgery (Sham *n* = 36, FLIT *n* = 18, equal numbers of females and males were used). **(a)** Representative trajectories of locomotion. Each tracing represents a travel trajectory of a mouse during a 10-minute testing period. **(b)** Percentage of time spent in central zone. **(c)** Latency of 1st entry into central zone (central zone: blue box in panel (a). Two-way ANOVA followed by post-hoc Bonferroni test was carried out to determine the *P* value of FLIT *vs*. sham. \*\*\**P* < 0.001; **** *P* < 0.0001. **(d to f)** Sexual behaviors test. **(d)** Mounting attempts was counted for male mice when female partners were present (*n* = 18 male per group). **(e)** Mounting attempts was counted for male mice when female partners were present (*n* = 18 male per group). **(f)** Urine testosterone levels of FLIT mice were significantly lower than the other groups at days 14 and 21 post surgery. Urine samples of three mice were pooled together which led to six urine samples per group for all time points. Two-way ANOVA followed by post-hoc Bonferroni test was carried out to determine the *P* value of FLIT *vs*. sham. \**P* < 0.05, \*\**P* < 0.01.

**Figure S4.**
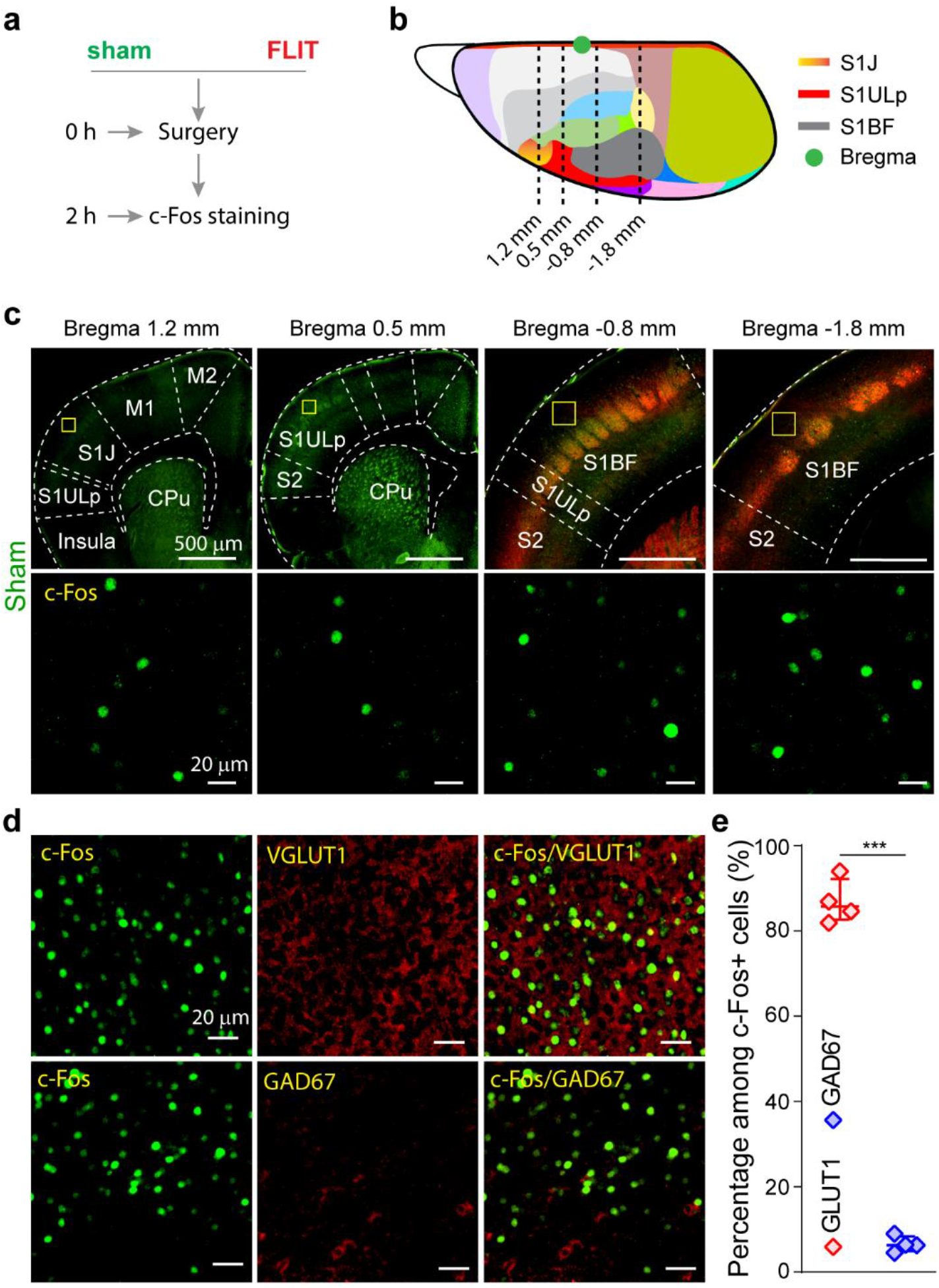
c-Fos expression pattern. **(a)** Diagram depicting c-Fos staining to assess neuronal activation after FLIT or sham surgery. **(b)** Schematic of mouse cortex with different colors representing specific cortical regions. Orange represents S1J, red represents S1ULp, and dark grey represents S1BF. Ten slices centered bregma 1.2 mm at 60 μm in thickness were used to assess c-Fos expression in S1J; fifteen slices centered bregma 0.5 mm at 60 μm in thickness were used to assess c-Fos expression in S1ULp; and fifteen slices between bregma −0.8 mm and −1.8 mm were used to assess c-Fos expression in S1BF. **(c)** Representative tangential slices of c-Fos staining of sham mice at 4X and 10X. Sequential slices from left to right represent coronal sections covering S1J (bregma 1.2mm), S1ULp (bregma 0.5mm), anterior S1BF (bregma −0.8mm) and posterior S1BF cortex (bregma −1.8mm). Slices between bregma - 0.8 mm and −1.8 mm were co-stained with VGLUT2 (red) to visualize barrels. Lower panels represent boxed regions of corresponding upper panels. **(d)** Representative staining of c-Fos and VGLUT1 or GAD67 in S1ULp-S1J of FLIT group. **(e)** Percentage of VGLUT1+ and GAD67+ cells among c-Fos+ cells. Two-tailed unpaired t-tests were carried out to determine the difference of GLUT1 *vs*. GAD67. Data are represented as mean ± SEM. *** *P* < 0.001.

**Figure S5.**
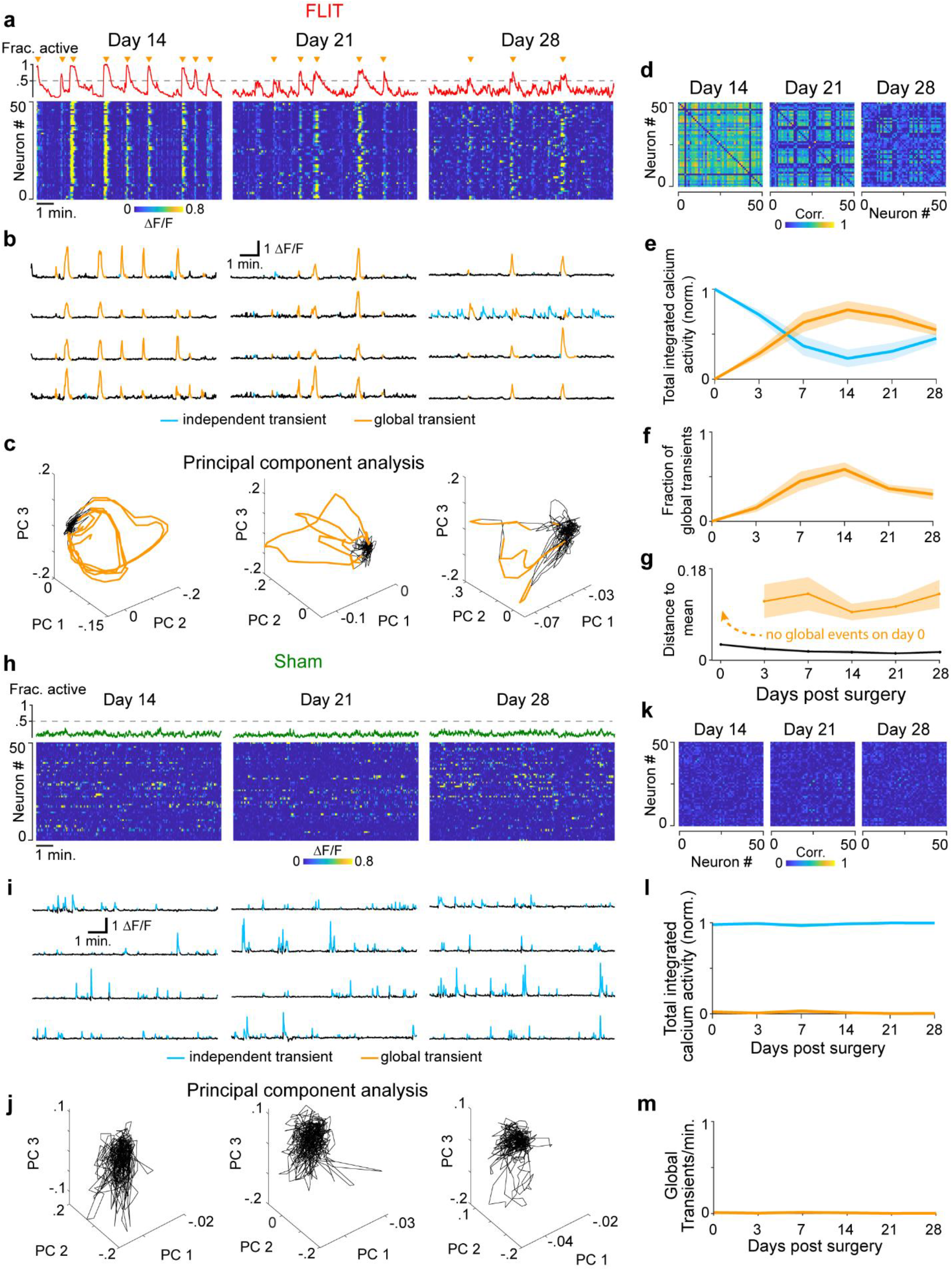
Neural activity on days 14-28. **(a)** Activity of all recorded neurons on days 14, 21 and 28 of the same FLIT mouse shown in Figure 2c. Fraction of simultaneously active neurons shown above heatmap. Global events are indicated by yellow triangles. **(b)** Four sample traces of calcium transients were shown with independent and global transients indicated. **(c)** Trajectories of neuronal populations plotted as a function of the first three principal components. Episodes highlighted in yellow correspond to global events. **(d)** Pairwise correlation matrices (Spearman’s rank correlation coefficient) for the days shown in (a). **(e)** Total integrated calcium activity for global and independent transients calculated as area under the curve (AUC), normalized to the total AUC. **(f)** Frequency of global transients as a function of days post FLIT surgery. Yellow lines indicate values for individual animals. **(g)** Distance of neuronal trajectory during global events to the mean for all FLIT animals. The distance was calculated as the Euclidean distance of the first three principal components from the mean value of those components. Indicated in yellow line is the mean Euclidean distance of datapoints belonging to global events. No global events were detected on day 0. **(h to m)** Same as above for the sham group.

**Figure S6.**
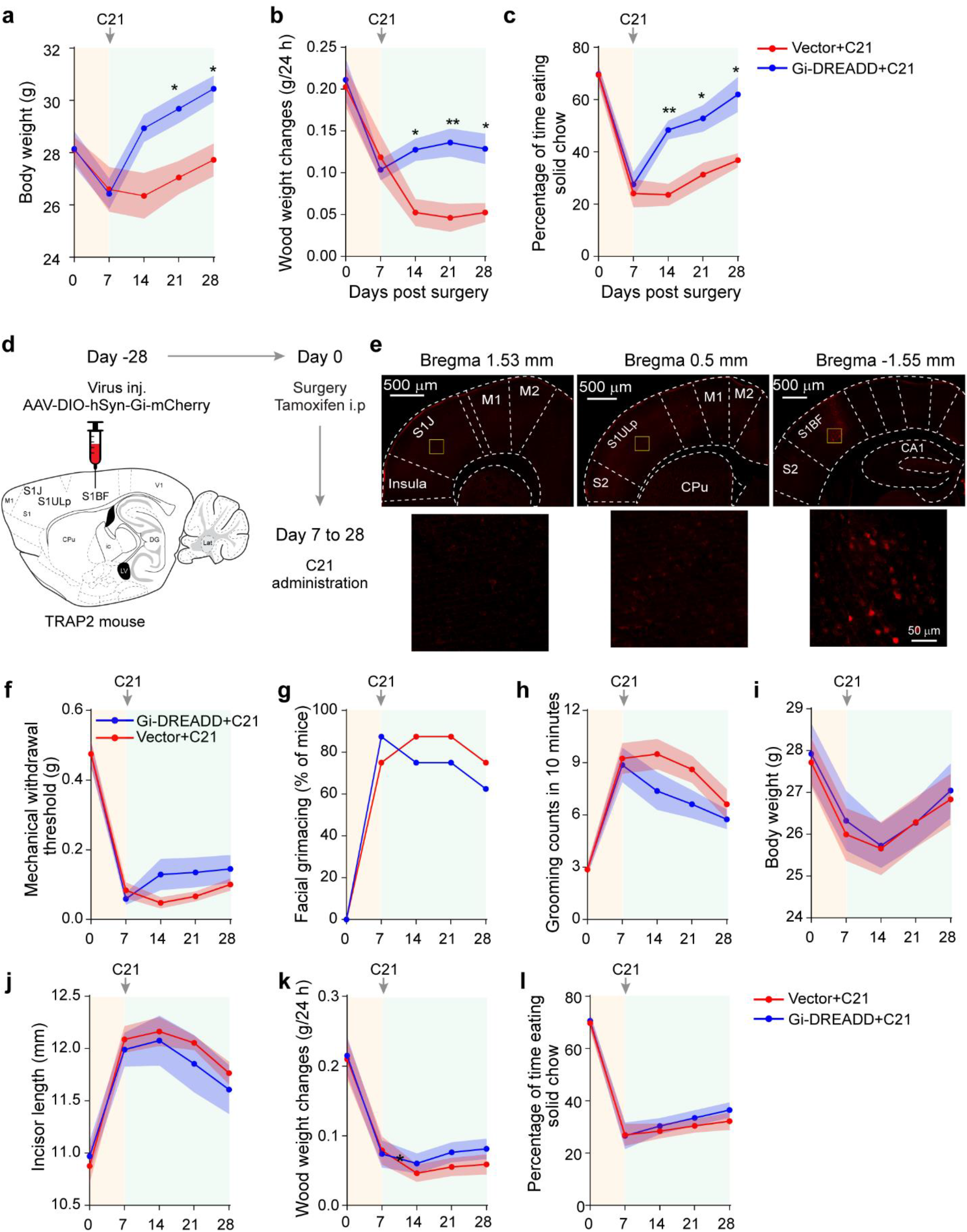
Chemogenetic manipulation of S1ULp-S1J but not S1BF suppresses pain-like behaviors. **(a to c)** Chemogenetic inhibition of c-Fos-induced Gi-expressing neurons leads to attenuated pain-like behavior. Behavioral testing was performed at indicated timepoints. At day 7, behavioral testing was performed prior to C21 administration to obtain pre-treatment baseline. **(a)** Body weight. **(b)** Wood weight changes in 24 hours. **(c)** percentage of time eating solid chow. Two-way ANOVA followed by Bonferroni post hoc test was carried out to determine the difference between the two groups. Data are represented as mean ± SEM. * *P* < 0.05, ** *P* < 0.01. **(d to l)** TRAP2 mouse was injected with AAV-DIO-hSyn-Gi-mCherry or AAV-DIO-mCherry vector in S1BF at day −28 (*n* = 8 per group). FLIT surgery was performed for all mice at day 0 accompanied by tamoxifen administration. From day 7, C21 was intraperitoneally administrated twice daily. Mice were sacrificed at day 28 for brain slices. **(d)** Diagram of virus injection and flowchart of experiment timeline. **(e)** Upper panels: Representative tangential slices of S1J, S1ULp and S1BF demonstrating mCherry expression was primarily located in S1BF. Lower panels represent boxed regions of corresponding upper panels. **(f)** mechanical withdrawal threshold to von Frey filaments. **(g)** Percentage of mice with facial grimacing. **(h)** Face grooming counts in 10 minutes. **(i)** Body weight. **(j)** Incisors length. **(k)** Wood weight changes in 24 hours. **(l)** Percentage of time eating solid chow. Two-way ANOVA followed by Bonferroni post hoc test was carried out to determine the difference between the two groups. Data are represented as mean ± SEM. Fisher’s exact test was used to determine statistical difference for facial grimaces.

**Figure S7.**
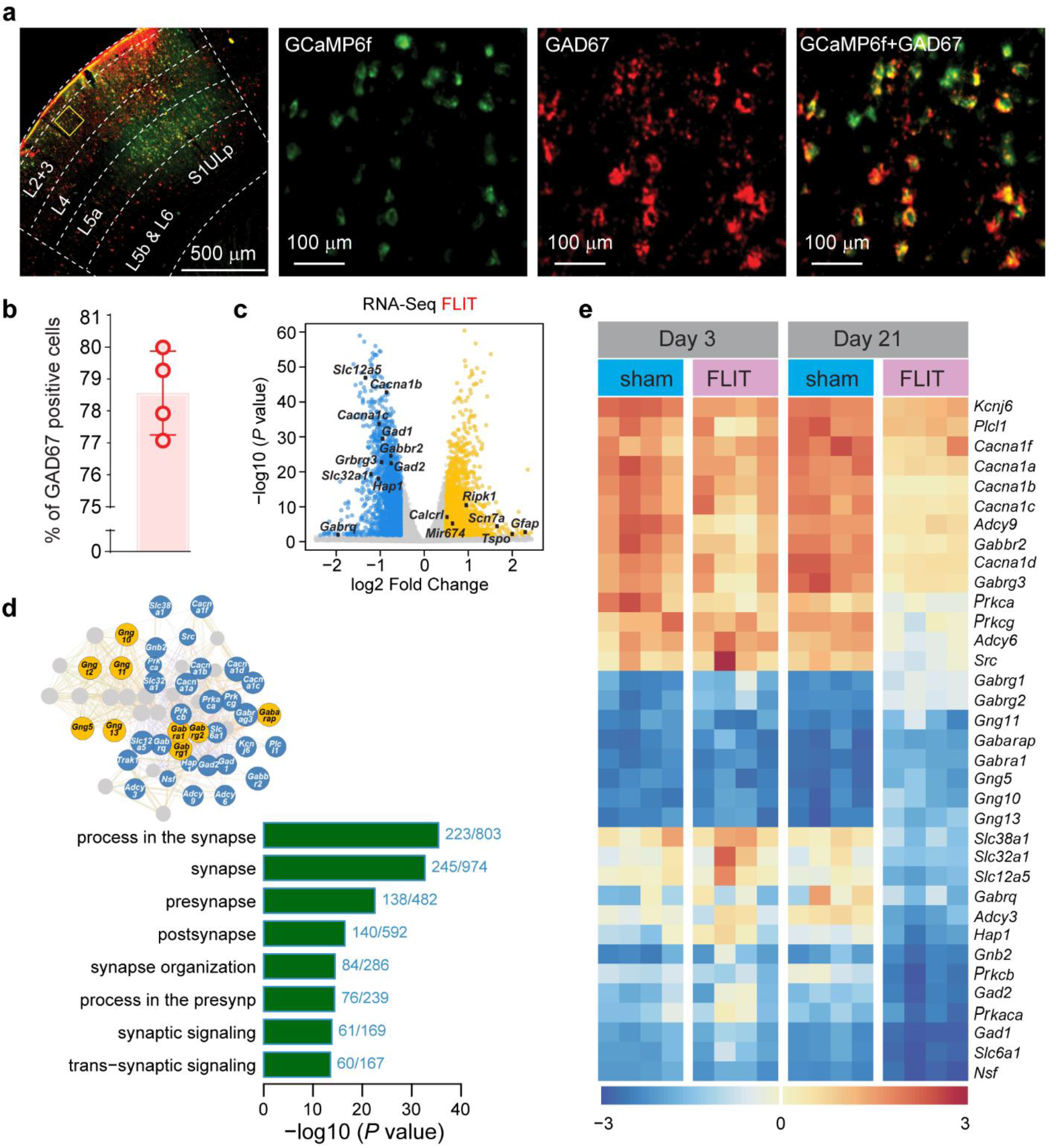
Cortical interneuron targeting and RNA-Seq of FLIT model. **(a and b)** AAV-Dlx-GCaMP6f expression in cortical interneurons. Mice were sacrificed 28 days after virus injection for brain slices. **(a)** Representative images of GAD67 staining (*n* = 4). **(b)** percentage of GAD67 positive cells among GCaMP6f expressing cells. **(c)** Interactome of differential regulated genes in FLIT model. Yellow: upregulated FLIT *vs*. sham; blue: downregulated FLIT *vs*. sham; cut-off threshold: 0.5 log2. **(d)** SynGO analysis of synapse-related pathway changes in the FLIT model. (**e)** Differentially regulated genes in the FLIT and sham mice (*n* = 4 each group, both normalized to Sham *n* = 4) at days 3 and 21 post surgery. Scale bar represents log2 fold changes.

**Figure S8.**
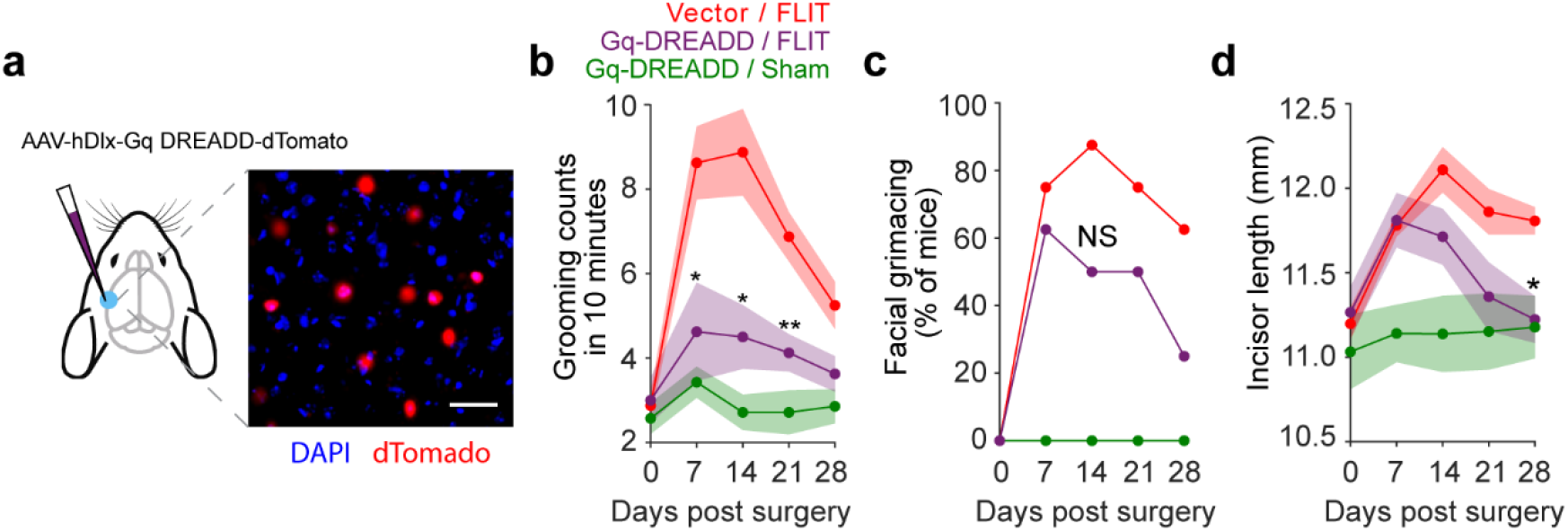
Chemogenetic manipulation of S1 interneurons. **(a)** Schematic and image of virus injection and expression (scale bar represents 50 μm). **(b)** Facial grooming counts in 10 minutes. Two-way ANOVA test indicates a significant difference among the groups. Post-hoc Bonferroni test was carried out to determine the *P* value of Vector / FLIT *vs*. Gq-DREADD / FLIT, \**P* < 0.05, \*\**P* < 0.01. **(c)** Percentage of mice with facial grimacing. Fisher exact test indicates no statistically significant difference among the groups. **(d)** Incisors length at indicated time points. Two-way ANOVA test indicates a significant difference among the groups. Post-hoc Bonferroni test was carried out to determine the *P* value of Vector / FLIT *vs*. Gq-DREADD / FLIT, \**P* < 0.05.

**Figure S9.**
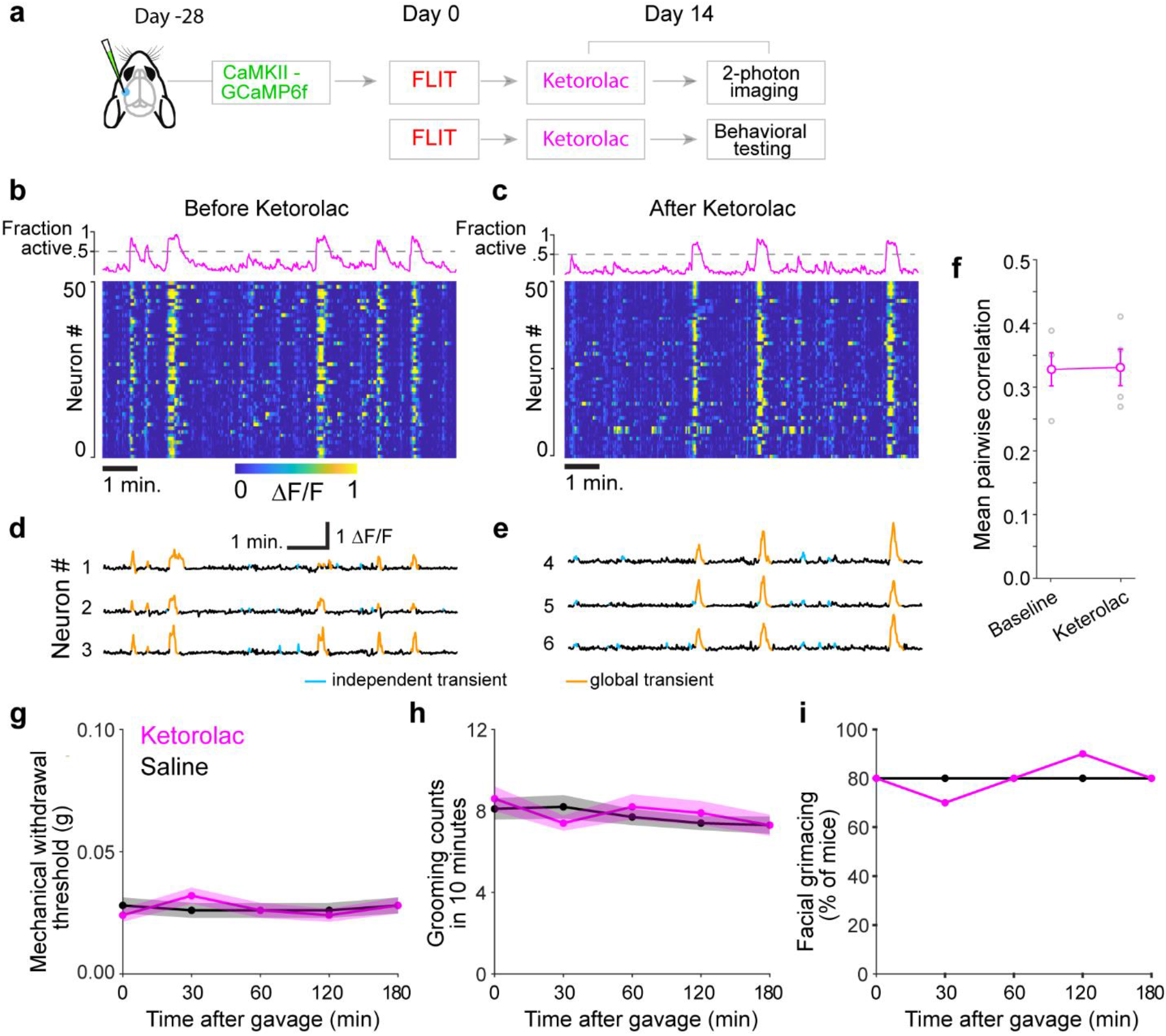
Ketorolac does not alleviate synchronized S1 activity or TN-like behaviors. Fourteen days post FLIT surgery, mice received ketorolac (10 mg/kg, oral gavage) treatment before excitatory neuron calcium imaging (*n* = 4) or behavioral testing (*n* = 10). **(a)** Flowchart of ketorolac experiment. **(b and c)** Representative heatmaps; fraction of active neuron plots; and **(d and e)** representative neuronal activity calcium transient traces in the same field of view before (panels B and D) and 60 minutes after (panels C and E) ketorolac administration. **(f)** Mean pairwise correlation before and after ketorolac treatment (Mean ± SEM: before: 0.33 ± 0.03, after: 0.33 ± 0.03, *P* = 0.93). **(g to i)** Ketorolac treatment does not alleviate TN-like behaviors in FLIT mice. **(g)** Mechanical withdrawal thresholds to von Frey filaments (mean ± SEM). **(h)** Facial grooming counts in 10 minutes at indicated time points. **(i)** Percentage of mice with facial grimaces.

**Figure S10.**
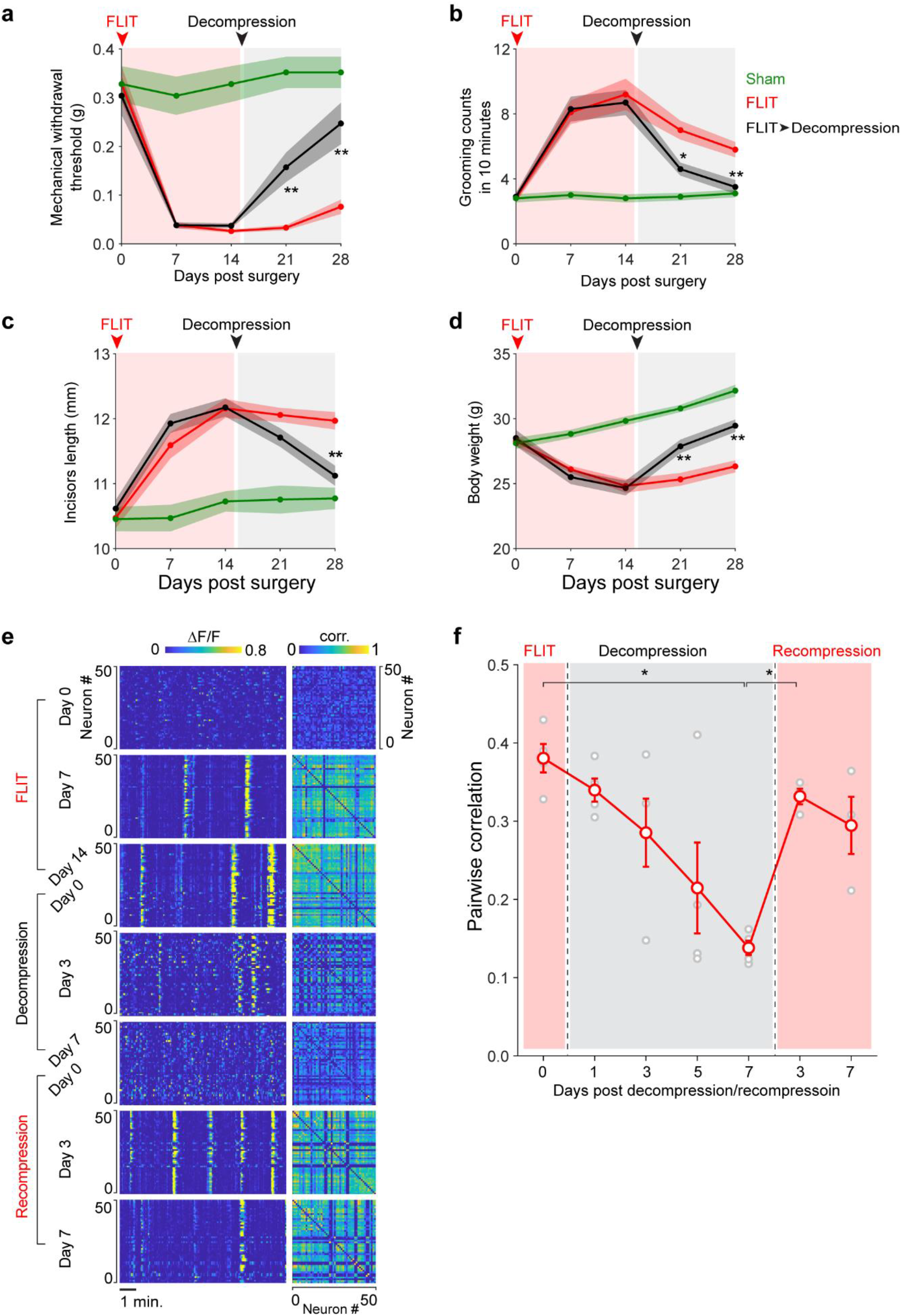
Nerve root decompression and recompression change cortical dynamics and pain-like behavior in FLIT mice. Fourteen days after FLIT surgery, mice (*n* = 10) received trigeminal nerve decompression. **(a)** Mechanical withdrawal; **(b)** Facial grooming counts in 10 minutes; **(c)** Incisors length; and **(d)** Body weight was compared among the groups. Two-way ANOVA test indicates significant difference was present among the groups, post-hoc Bonferroni test was carried out to determine the *P* value of FLIT *vs*. FLIT + decompression \**P* < 0.05; \*\**P* < 0.01. **(e and f)** Reoccurrence of S1 synchronization is induced by recompression of trigeminal nerve root in decompressed FLIT mice (*n* = 3). Fourteen days post FLIT surgery, mice received decompression by removing Surgifoam through foramen lacerum. Seven days after decompression, mice underwent recompression of trigeminal nerve root. **(e)** Representative S1 excitatory neuronal activity heatmaps and correlation matrices at indicated time points. **(f)** Pairwise correlation of neuronal activity over the time course of the decompression and recompression experiment (One-way ANOVA: *P* < 0.05, Turkey-Kramer post hoc comparison significant for between day 7 decompression and day 3 recompression), \**P* < 0.05.

